# Circadian rhythms and the light-dark cycle interact to regulate amyloid plaque accumulation and tau phosphorylation in 5xFAD mice

**DOI:** 10.1101/2025.03.31.645805

**Authors:** Melvin W. King, Sophie Jacob, Ashish Sharma, Jennifer H. Lawrence, Erik S. Musiek

**Affiliations:** Department of Neurology & Center On Biological Rhythms & Sleep, Washington University School of Medicine in St. Louis, St. Louis, MO, USA

**Keywords:** circadian rhythms, Alzheimer’s Disease, amyloid, suprachiasmatic nucleus, tau

## Abstract

**Background:** Circadian disruption has long been appreciated as a downstream consequence of Alzheimer’s Disease in humans. However, an upstream role for behavioral circadian disruption in regulating AD pathology remains an open question.

**Methods:** To determine the role of the central circadian clock in the suprachiasmatic nucleus (SCN) in regulating amyloid pathology, we generated mice harboring deletion of the critical clock gene *Bmal1* in GABAergic neurons using VGAT-iCre, which is expressed in >95% of SCN cells, and crossed this line to the 5xFAD amyloid mouse model. To examine the role of the light-dark cycle in this process, we aged these mice in either regular 12:12 light-dark (LD) or constant darkness (DD) conditions. Transcriptional, behavioral, and physiological rhythms were examined in VGAT-iCre; *Bmal1*^fl/fl^; 5xFAD (VGAT-BMAL1KO;5xFAD) mice under varying light conditions. Amyloid plaque deposition, peri-plaque tau phosphorylation, glial activation, and transcriptomic changes were examined.

**Results:** VGAT-BMAL1KO;5xFAD mice showed loss of SCN BMAL1 expression and severe disruption of behavioral rhythms in both LD and DD, with loss of day-night rhythms in consolidated sleep and blunting of rhythmic clock gene expression in the brain. Surprisingly, VGAT-BMAL1KO;5xFAD mice kept under LD showed reduced total plaque accumulation and peri-plaque tau phosphorylation, compared to Cre-negative controls. These changes were gated by the light-dark cycle, as they were absent in VGAT-BMAL1KO;5xFAD mice kept in DD conditions. Total plaque accumulation was also reduced in control 5xFAD mice kept in DD as compared to LD, suggesting a general effect of light-dark cycle on amyloid aggregation. Expression of murine presenilin 1 (*Psen1*), as well as amyloidogenic cleavage of amyloid precursor protein, were suppressed in VGAT-BMAL1KO;5xFAD specifically under LD conditions.

**Conclusions:** In 5xFAD mice, the central circadian clock and the light-dark cycle interact to regulate amyloid pathology. Disruption of the central clock in the presence of a light-dark cycle may reduce APP cleavage and plaque formation. These results call into question the proposed simple positive feedback loop between circadian rhythm disruption and Alzheimer’s Disease pathology.

## Introduction

Alzheimer’s Disease is a progressive, debilitating neurological disorder that effects nearly 6.9 million people aged 65 or over and is the leading cause of dementia in the United States ^1^. Disruption of endogenous circadian and sleep rhythms has long been appreciated as a downstream consequence of Alzheimer’s Disease ^2,3^. However, an upstream role for circadian disruption in AD progression -- and in particular the aggregation of Aβ plaques -- has yet to be defined ^4^. Evidence consistent with a causal role for circadian disruption comes from numerous human studies demonstrating transcriptional and behavioral circadian abnormalities as a function of age, the primary risk factor for developing AD ^5,6^. The presence of preclinical Aβ or tau pathology in cognitively normal individuals has been associated with increased circadian fragmentation, suggesting that circadian dysfunction occurs very early in the disease course ^7^. Given the pervasiveness of the circadian system at various levels of biology, how these disruptions contribute to the initiation and progression of AD pathology is a significant unknown.

A few previous studies have examined the impact of circadian disruption on AD pathology in mouse models using genetic ablation of the transcription factor *Bmal1* (aka *Arntl*), as BMAL1 protein dimerizes with CLOCK or NPAS2 to drive cellular rhythms through the core circadian Transcriptional-Translation Feedback Loop (TTFL)^8,9^. Our group previously found that post-natal whole-body knockout of *Bmal1* causes behavioral arrhythmicity in mice and exacerbates hippocampal plaque pathology in the APP/PS1- 21 model of AD^10,11^. However, a suprachiasmatic nucleus (SCN)-sparing *Bmal1* deletion failed to have the same impact on plaque deposition ^12^. This suggests that disrupting *Bmal1* across cell types in the brain and periphery may exert complicated circadian and non-circadian effects that impact amyloid dynamics. It remains unknown whether specifically disrupting rhythmic output from the SCN, which coordinates brain- and body-wide rhythms by a combination of synaptic and hormonal cues and whose rhythmic output degrades with age, may influence Aβ plaque pathology ^13,14^.

One classic strategy for disrupting SCN function is through electrolytic lesion^15^. However, the SCN is small and surrounded by dozens of other critical hypothalamic nuclei which may be variably damaged in this process, exerting unknown effects^16^. Thus, a genetic approach is preferable. Since more than 95% of SCN neurons are GABAergic, targeted deletion of *Bmal1* in GABAergic neurons using VGAT-ires-Cre produces mice which are behaviorally arrhythmic in constant darkness (DD) ^17,18^. It is unknown how this restricted *Bmal1* deletion impacts amyloid pathology in mice. However, mice with a genetically ablated clock do maintain some rhythmic behavior under regular light-dark lighting cycles due to phenomenon called masking, and the effect of this on amyloid pathology is also unknown^19,20^.

Here, we have examined the effect of GABAergic *Bmal1* deletion on circadian function and amyloid-related pathology in the 5xFAD mouse model of amyloid pathology^21^. We performed experiments with VGAT-iCre; *Bmal1*^fl/fl^;5xFAD^+/-^ mice (referred to as VGAT- BMAL1KO;5xFAD or 5x;Cre+) and Cre-;*Bmal1*^fl/fl^;5xFAD^+/-^ (5x;Cre-) controls under both standard 12h:12h light:dark (LD, in which Cre+ mice have some rhythmic behavior due to zeitgeber cues) and DD conditions (under which Cre+ mice are fully arrhythmic) to help isolate the potential effects of the light-dark cycle and circadian rhythmicity from non-circadian effects of *Bmal1* deletion. To our surprise, we have found that disruption of behavioral and transcriptional rhythms via GABAergic *Bmal1* deletion caused a decrease in amyloid plaque accumulation and plaque-related tau phosphorylation, but only in mice kept under LD conditions. These studies suggest that specific interactions between the light-dark cycle and the molecular clock may drive changes in Aβ aggregation, and provoke a rethinking of proposed linear relationship between behavioral arrhythmicity and AD pathology.

## Results

### GABAergic *Bmal1* deletion disrupts circadian behavior in light-dark and constant darkness

We sought to use the VGAT-ires-Cre line to target clock gene expression in the SCN, as previously described, so as to disrupt SCN circadian function without widespread deletion of clock genes in other cell types and regions of the brain^17^. Inhibitory GABAergic synaptic transmission is ubiquitous throughout the brain, so we first aimed to characterize the expression of VGAT-ires-Cre in various brain regions. We crossed VGAT-iCre mice to the Ai9 mouse line expressing tdTomato upstream of a *loxP*-flanked STOP cassette, enabling permanent labeling of VGAT-expressing cells ^22^. We observed strong tdTomato labeling within the SCN (Figs 1A, S1A), in agreement with previous reports ^23^. Intense tdTomato signal was noted in the striatum and basal ganglia (Fig. S1A), as most of these neurons are GABAergic. However, in the medial cortex and hippocampus, regions typically examined for plaque accumulation in AD models, only sparse labeling was seen with roughly 20% of NeuN+ expressing tdTomato across both regions (Fig. S1B).

**Figure 1.**
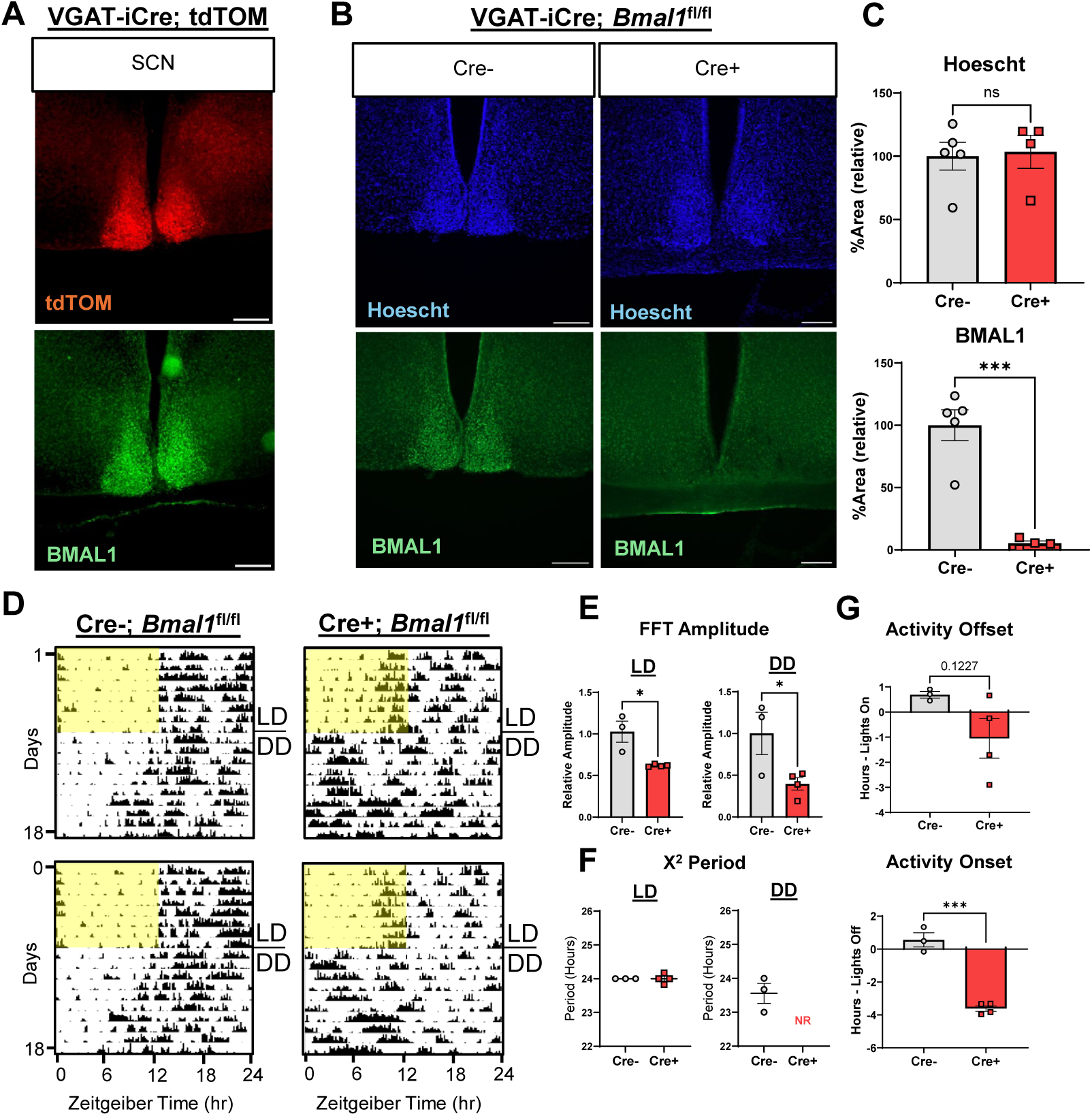
Disruption of circadian behavior in light-dark and constant darkness by GABAergic *Bmal1* deletion. **A.** Immunostaining of BMAL1 (green) with auto-fluorescent tdTomato (red) on SCN sections of *VGAT-ires-iCre;tdTOM* lox-stop-lox (Ai9) mice. Scale bars are 200 µm **B.** Double immunostaining of DAPI (blue) and BMAL1 (green) on SCN sections of *VGAT-ires-Cre;Bmal^fl/fl^* mice. Scale bars are 200 µm **C.** Quantification of staining as shown in (B) (Cre+: *VGAT-ires-iCre+;Bmal1^fl/fl^*, n=5, Cre-: *Bmal1*^fl/fl^, n=4) values are mean +/- SE (*** Student’s t, p < 0.0005) **D**. Representative actograms of *VGAT-ires-iCre;Bmal1^fl/f^* mice. All mice were placed into 12:12 LD for at least 7 days before being transferred into DD, as indicated on the plot. Yellow bars represent days of LD -- hour 6 to hour 18.. **E.** Quantification of FFT Amplitude from actogram data plotted in (D). LD analysis was performed on data generated on days 2-9. DD analysis data was taken from days 12-19. *VGAT-ires-iCre+;Bmal^fl/fl^* (n = 3-4 mice per genotype as indicated; *Student’s t, p < 0.05) **F.** Χ^2^ periodogram analysis from actogram data plotted in (D). LD period analysis was performed on data generated on days 2-9. DD analysis data was taken from days 12-19. NR = No Coherent Rhythms. **G.** (Top) Average activity offset time (see methods) minus lights on time (ZT0 or hour 6). (Top) Average activity onset time minus lights off (ZT12 or hour 18) (n=3-4 mice per group as indicated *** Student’s t, p < 0.0005).

Given the role of the suprachiasmatic nucleus and BMAL1 in maintaining organismal and cellular circadian rhythms, we then assessed the role of GABAergic *Bmal1* in regulating circadian behavior under light-dark and constant darkness. We crossed VGAT-ires-Cre and *Bmal1*^fl/fl^ mice to generate Cre-; *Bmal1*^fl/fl^ controls and Cre+ VGAT- BMAL1 KO mice and aged them to two months. Circadian activity was assessed with infrared actigraphy monitoring under LD (1 week) and DD (2 weeks) conditions. Under LD conditions, Cre+ mice showed some diurnal rhythms in activity but had noticeably dampened amplitude (Fig 1D,E), and had onset of their active phase nearly four hours earlier than Cre- mice, which normally become active at lights off (ZT12, Fig 1G). Upon release into constant darkness, Cre- mice maintained circadian activity rhythms with an average period of 23.6 hours, while Cre+ mice immediately lost coherent rhythmic activity by periodogram analysis, with no detectable circadian period (Fig 1F, S1D). BMAL1 immunohistochemical (IHC) analysis confirmed the near total loss of BMAL1 protein within the SCN (Fig 1B, 1C). Interestingly, despite intense tdTomato expression within the striatum, VGAT-iCre mediated excision only reduced BMAL1 protein expression within this region by roughly half, whereas total BMAL1 protein expression at the IHC level within the cortex and hippocampus was unaffected, suggesting that Bmal1 deletion is confined to the SCN and striatum, and that GABAergic interneurons are not major expressors of *Bmal1* in the cortex and hippocampus (Fig S1C). Collectively, these results confirm that GABAergic *Bmal1* deletion disrupts SCN circadian function with minimal impact on BMAL1 expression in cortex and hippocampus.

### GABAergic *Bmal1* deletion disrupts sleep and transcriptional rhythms in 5xFAD mice

Our next goal was to explore how GABAergic BMAL1 influences rhythmic physiology in a mouse model of Alzheimer’s Disease. We crossed 5xFAD mice, which express human amyloid precursor protein (APP) with the Swedish (KM670/671NL), Florida (I716V), and London (V717I) mutations and human presenilin 1(PSEN1) with the M146L and L286V mutations, to the VGAT-ires-Cre;*Bmal1*^fl/fl^ line to generate VGAT-iCre; *Bmal1*^fl/fl^;5xFAD^+/-^ mice (referred to as VGAT-BMAL1KO;5xFAD or 5x;Cre+) and Cre-; *Bmal1*^fl/fl^;5xFAD^+/-^ controls (5x;Cre-) (Fig 2A)^21^. Mice were aged in LD until 5 months, when substantial plaque development has taken place within the cortex and hippocampal regions^24^. As before, while 5x;Cre+ mice were able to entrain to a regular LD cycle, their circadian amplitude was negatively impacted compared to 5x;Cre- (Fig 2B, 2C, S2A). Constant darkness abrogated rhythms in 4/4 Cre+ mice by χ^2^ periodogram analysis, demonstrating the efficacy of this paradigm in ablating rhythms in 5xFAD mice in constant darkness (Fig. S2B).

**Figure 2.**
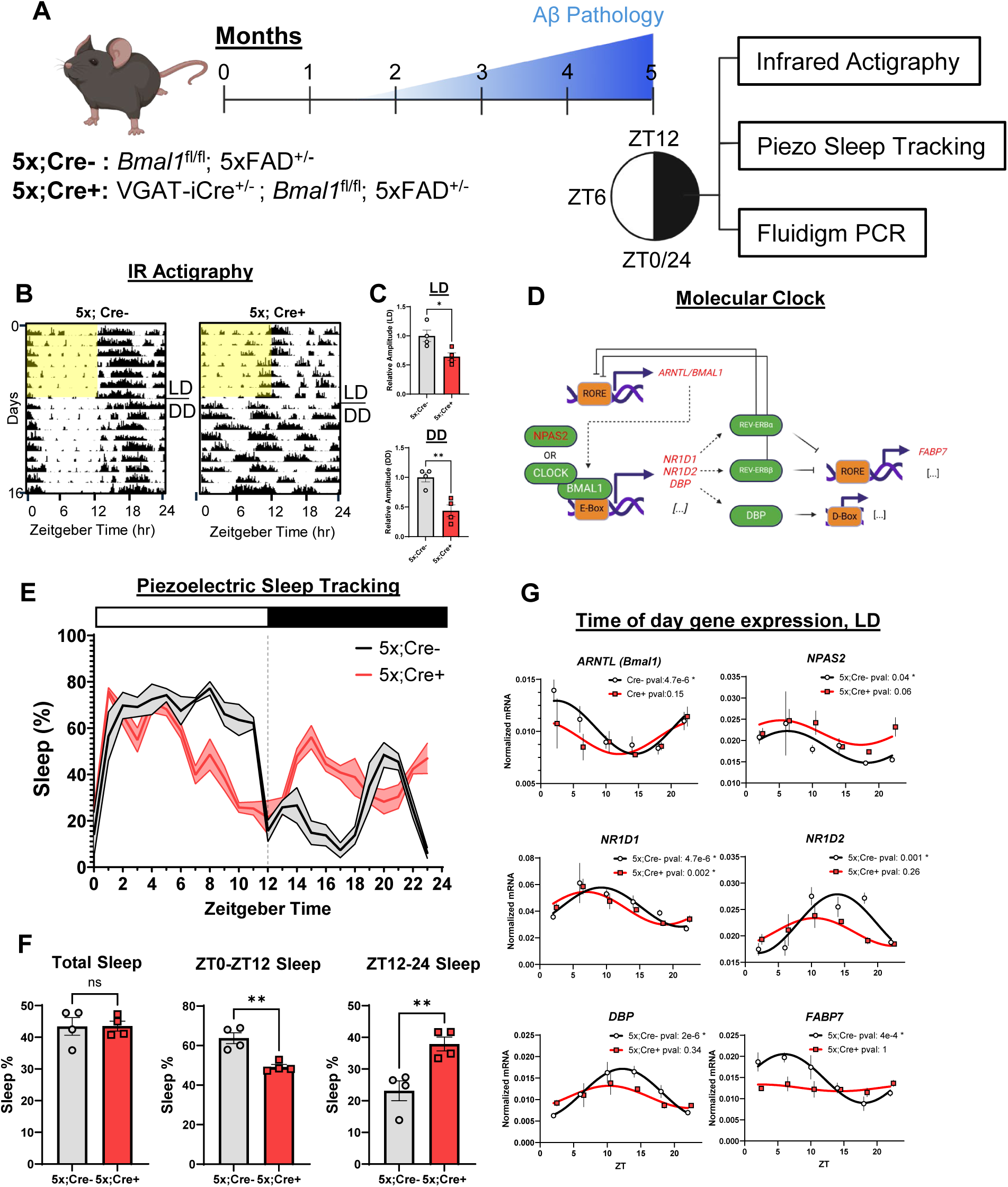
GABAergic *Bmal1* deletion disrupts behavioral, sleep, and transcriptional rhythms in 5xFAD mice. **A.** Schematic of the experimental paradigm used to assess physiology in Cre+ *VGAT-ires- Cre;Bmal^fl/fl^;*5xFAD mice (VGAT-BMAL1KO;5xFAD or 5x;Cre+) and Cre- *Bmal^fl/fl^;*5xFAD (5x;Cre-) littermate controls. **B.** Representative actograms from 5x;Cre+ and 5xFAD;Cre- mice. Mice were housed in 7 days of LD before being released into DD. **C.** Quantification of amplitude from actograms plotted in B (n=4 mice per group as indicated, *p < 0.05 or ** p < 0.005 by Student’s t) **D.** Diagram of the molecular clock, with measured genes highlighted in red. **E.** Rolling hourly average sleep percentage in 5x;Cre+ mice and 5x;Cre- controls as a function of time. Black trace: Cre-negative. Red trace: Cre-positive. Zeitgeber time ZT0 defined as the start of the light period. **F.** Quantification of Total, ZT0-12 and ZT12-24 Sleep percentage. (**p < 0.005 by Student’s t) **G.** Data points and cosinor fit of selected clock genes. Black circles are Cre-, Red squares are Cre+ (n=2-5 mice per genotype per time point. Benjamin-Hochberg adjusted p-values are displayed above each graph).

We simultaneously measured sleep -- a circadian controlled process whose disruption is associated with risk of dementia in humans -- using piezoelectric monitoring ^25,26^. Total sleep percentage over a 24-hour period was unchanged between 5x;Cre+ and 5x;Cre- control mice. However, in accordance with their earlier onset time, 5x;Cre+ mice spent less time sleeping during the inactive or light phase, particularly from ZT6 to ZT12 (Fig 2E, 2F). By comparison, 5x;Cre+ mice spent a larger percentage of time sleeping during the dark phase, particularly between ZT14 and ZT18. Free-running behavior (5x;Cre-) and circadian fragmentation (5x;Cre+) prevented division of sleep behavior in DD into active and inactive epochs, but overall sleep amount remained the same between Cre- and Cre+ mice, similar to LD (Fig S2C). Despite significant circadian disruption between 5x;Cre- and 5x;Cre+ mice, the total amount of sleep, to which plaque deposition in mouse models is strongly sensitive, remained unchanged ^27–29^.

Finally, we examined the role of GABAergic BMAL1 in driving hippocampal diurnal transcriptional rhythms under LD conditions. Bulk hippocampal tissue was harvested from 5x;Cre+ mice and 5x;Cre- controls across six times of day under LD, and analyzed for a panel of transcripts associated with circadian and clock function (Fig 2D). Because 5x;Cre+ mice are arrhythmic in DD, we did not perform gene expression under these conditions, as the phase dispersion of the Cre+ mice at each timepoint would make the data uninterpretable. We examined eight known circadian transcripts by qPCR, and all were identified as circadian via JTK_CYCLE analysis in 5x;Cre- control mice under LD ^30^. However, all showed blunted amplitude in 5x;Cre+ mice under LD, with only *Nr1d1* (encoding Rev-Erbα) remaining rhythmic, although phase advanced by 2 hours (Fig 2G, S2C). Thus, despite having partially intact behavioral rhythms under LD conditions, circadian gene expression is disrupted in the hippocampus of 5x;Cre+ mice. Together, our data demonstrate that GABAergic *Bmal1* KO in 5xFAD mice disrupts both LD and DD circadian behavior, disrupts sleep timing and dampens oscillatory hippocampal clock gene expression.

### GABAergic *Bmal1* knockout suppresses amyloid plaque pathology in light-dark

As circadian disruption is a hallmark of preclinical AD, we next tested the hypothesis that excision of *Bmal1* in GABAergic neurons might influence Aβ aggregation in 5xFAD mice^7^. We used exclusively female VGAT-BMAL1KO;5xFAD (5x; Cre+) mice and 5x;Cre- littermates, as females have more aggressive plaque pathology. We kept mice under standard LD conditions until 5 months of age, when plaque accumulation is moderate^24^. At 5 months, staining with anti-Aβ antibody HJ3.4, which labels total Aβ plaque (diffuse and fibrillar), revealed a surprising reduction in plaque density in 5x;Cre+ mice within the cortex of roughly 40%, with smaller reductions of 25% in the hippocampal and thalamic regions (Fig 3A, 3B). As a partial explanation for these changes, we found a mild (∼10%) but statistically significant reduction in plaque size within the cortex and hippocampus, along with a trending but non-significant reduction in plaque numbers in these regions (Fig 3D, 3E). Looking specifically at plaque numbers, histogram analysis revealed a significant reduction in the number of smaller plaques, between 30 and 250 square microns, in both the cortex and hippocampus (Fig 3F).

**Figure 3.**
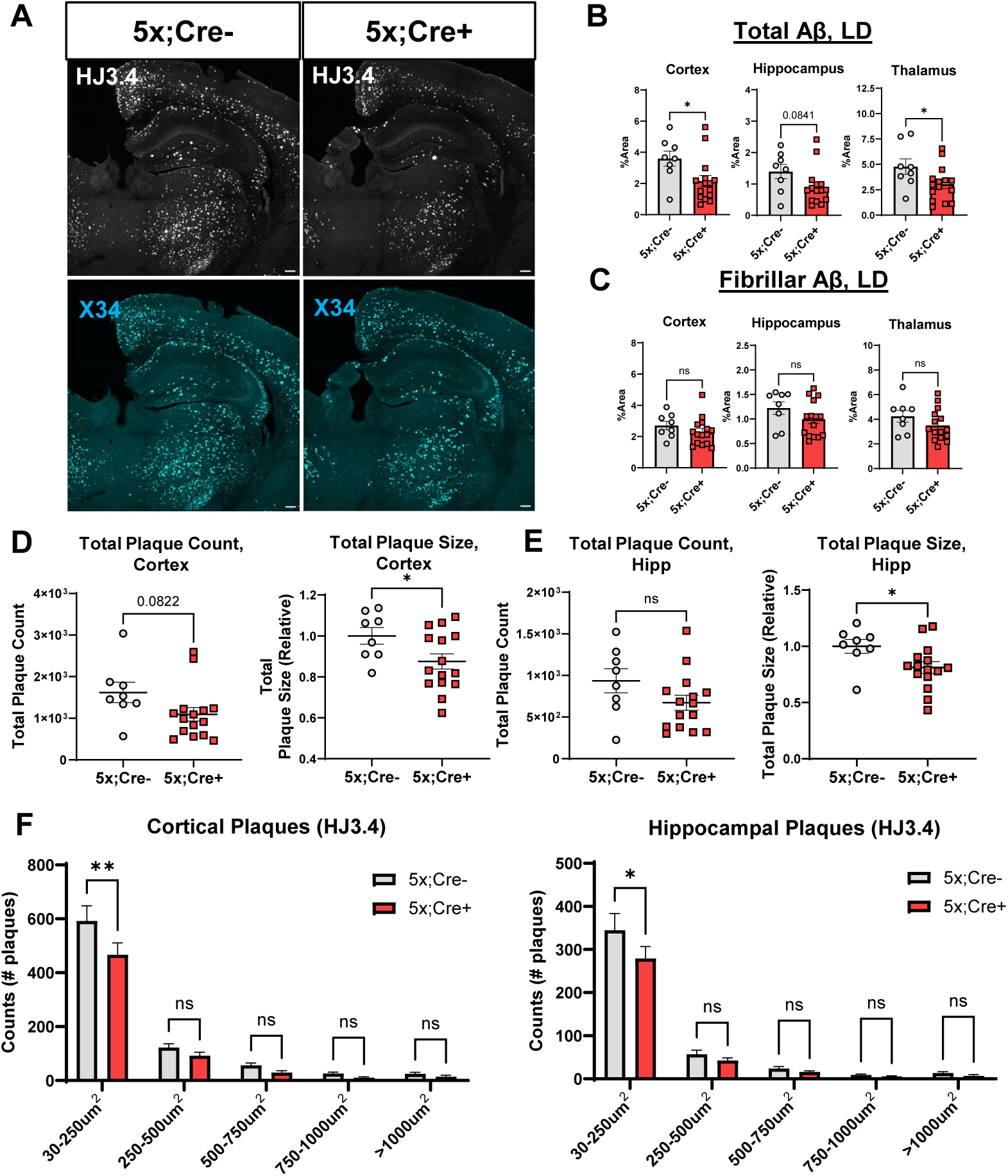
GABAergic *Bmal1*-knockout mitigates plaque accumulation under light-dark conditions. **A.** Representative whole brain images from 5mo 5x;Cre+ mice and Cre-; 5xFAD control mice aged in LD, depicting staining by X34 (fibrillar plaques, cyan) and HJ3.4 (total plaques, white). (Scale bar = 200 µm) **B-C.** Quantification of regional plaque %area as depicted in A. (n=9-15 mice per group as indicated. ns = not statistically significant, *p<0.05 by Students t, p-values between 0.05 and 0.1 are displayed) **D-E.** Quantification of cortical and hippocampal plaque total (HJ3.4) counts (first panel) and size (second panel) (n = 9-15 mice per group as indicated. *p < 0.05 by Students t, p-values between 0.05 and 0.1 are displayed) **F.** Histogram of total (HJ3.4) plaque counts in the cortex (left panel) and hippocampus (right panel) (n=9-15 mice per group, *p<0.05 or **p<0.05 by Two-Way ANOVA with Šídák’s multiple comparisons test)

We next examined the accumulation of dense core fibrillar Aβ plaque using the congophilic stain X34. Despite no change in total X34 accumulation in VGAT- BMAL1KO;5xFAD mice across regions, confocal imaging revealed changes in fibrillar plaque morphology typically indicative of lower toxicity: fibrillar plaques in the hippocampus in particular were more compact, with a smaller size and higher circularity index (Fig S3D)^31^. We co-stained sections with X34 and HJ3.4b and found volumetric reductions in HJ3.4+ halo of diffuse amyloid around the volume of fibrillar X34+ plaques, as well as a reduction in the amount of diffuse, non-X34 associated HJ3.4, particularly within the hippocampus (Fig 3B, 3C). These data suggest that GABAergic *Bmal1* knockout alters the morphology of Aβ plaques and mitigates the accumulation of non-fibrillar amyloid.

### GABAergic *Bmal1* knockout mitigates neuropil tau accumulation in light-dark

We next examined peri-plaque pathological changes in VGAT-BMAL1KO;5xFAD mice and Cre-; 5xFAD control mice. Growth of amyloid plaques induces a combination of maladaptive responses in nearby neurites including axonal swelling, overexpression of Aβ-cleavage and lysosomal enzymes, and promotion of tau phosphorylation ^32–34^. We stained for dystrophic neurites using an antibody against the lysosomal enzyme LAMP1 and colocalized them around X34 plaques (Fig 4A). We found no major change in the total volume of neuritic dystrophy nor in the amount of dystrophy normalized per X34 plaque (Fig 4A, 4B) between genotypes in any of the regions imaged. However, when we stained for neuropil (NP) phosphorylated (S202 & T205) tau using the antibody AT8 we found a dramatic reduction in VGAT-BMAL1KO;5xFAD mice as compared to 5x;Cre- controls, with more than 75% of AT8 in the vicinity of imaged X34 plaques reduced per unit of X34 volume in both the hippocampus and thalamus (Fig 4C, 4D).

**Figure 4.**
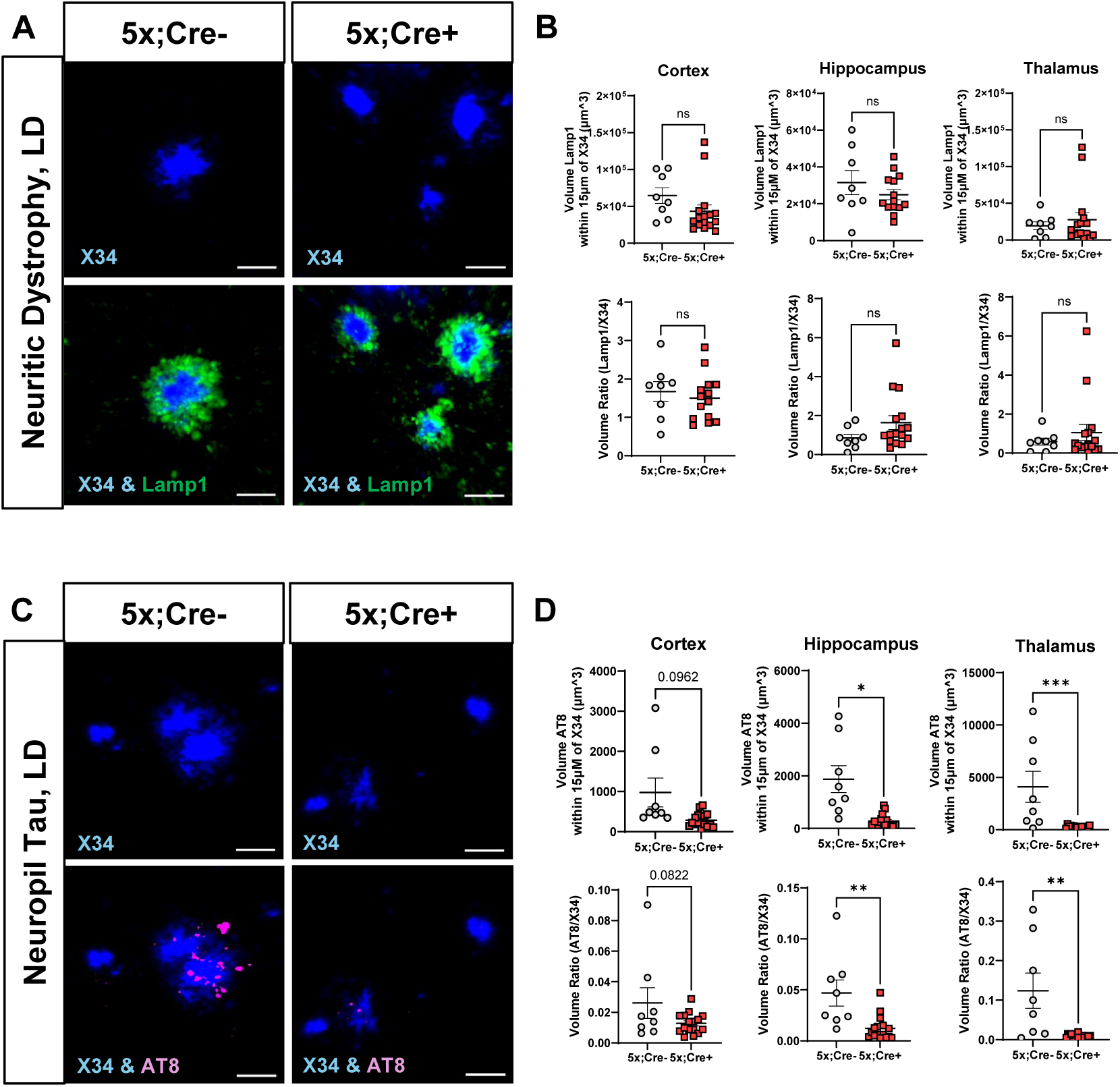
Diminished neuropil tau aggregation in GABAergic *Bmal1* KO mice under light-dark conditions. **A.** Representative confocal images depicting fibrillar plaque (blue, X34) and neuritic dystrophy (green, LAMP1) in 5mo 5x;Cre+ mice and 5x;Cre- control mice aged in LD (Scale bar, 20 µm). **B.** Quantification of LAMP1 as depicted in (A) Top: averaged total LAMP1 volume per field. Bottom: total LAMP1 volume per field divided by X34 volume. (n = 9-15 mice per group, ns=not statistically significant **C.** Representative confocal images depicting fibrillar plaque (blue, X34) and neuropil tau (magenta, AT8) (Scale bar, 20 µm) in 5mo 5x;Cre+ mice and 5x;Cre- controls aged in LD. **D.** Quantification of AT8 as depicted in (C) Top: averaged total AT8 volume per field. Bottom: total AT8 volume per field divided by X34 volume.) (n = 9-15 mice per group, *p < 0.05 or **p<0.005 or ***p<0.0005 by Students t, p-values between 0.05 and 0.1 are displayed)

We next examined microglia, brain parenchymal immune cells who are capable of both protective and detrimental functions in the presence of Aβ pathology, to determine if they might be implicated in the amyloid and tau changes seen in VGAT- BMAL1KO;5xFAD mice ^35,36^. We used confocal imaging to characterize Iba1/CD68 double-positive microglia around X34+ plaques, which indicate phagocytic activated microglia. Though a trending decrease in Iba1+ volume around plaques in the cortex was noted, we saw no other major changes in any of our microglia markers between VGAT-BMAL1KO;5xFAD mice and 5x;Cre- mice (Fig S4A, S4B). We also performed transcript analysis for various homeostatic and disease-associated microglia (DAM) markers in bulk hippocampal tissue and saw no changes (Fig S4C) ^37^. These data suggest that GABAergic *Bmal1* knockout in the setting of light-dark reduces peri-plaque phospho-tau aggregation, without changing neuritic dystrophy or Aβ-induced microgliosis.

### GABAergic *Bmal1* knockout has minimal effect on plaque pathology in constant darkness

The effect of GABAergic *Bmal1* knockout on circadian behavior can be partially masked by the light-dark cycle, so we next asked if allowing Cre+ mice to be fully arrhythmic might modulate the effect on amyloid pathology. Starting at six weeks of age, we placed VGAT-BMAL1KO;5xFAD (and 5x;Cre- littermates, all female) in constant darkness until age 5 months, when they were harvested and analyzed immunohistochemically for Aβ aggregation and related pathology as above. Surprisingly, VGAT-BMAL1KO;5xFAD mice kept in DD displayed no differences in total or fibrillar plaque density in any of the analyzed regions, as compared to 5x;Cre- littermate controls (Fig 5A, 5B). No effect was seen on peri-plaque NP-tau aggregation using AT8. However, VGAT-BMAL1KO;5xFAD mice did show a consistent trend toward reductions in LAMP1+ neuritic dystrophy in all three regions, though none reached individual statistical significance (Fig 5D).

**Figure 5.**
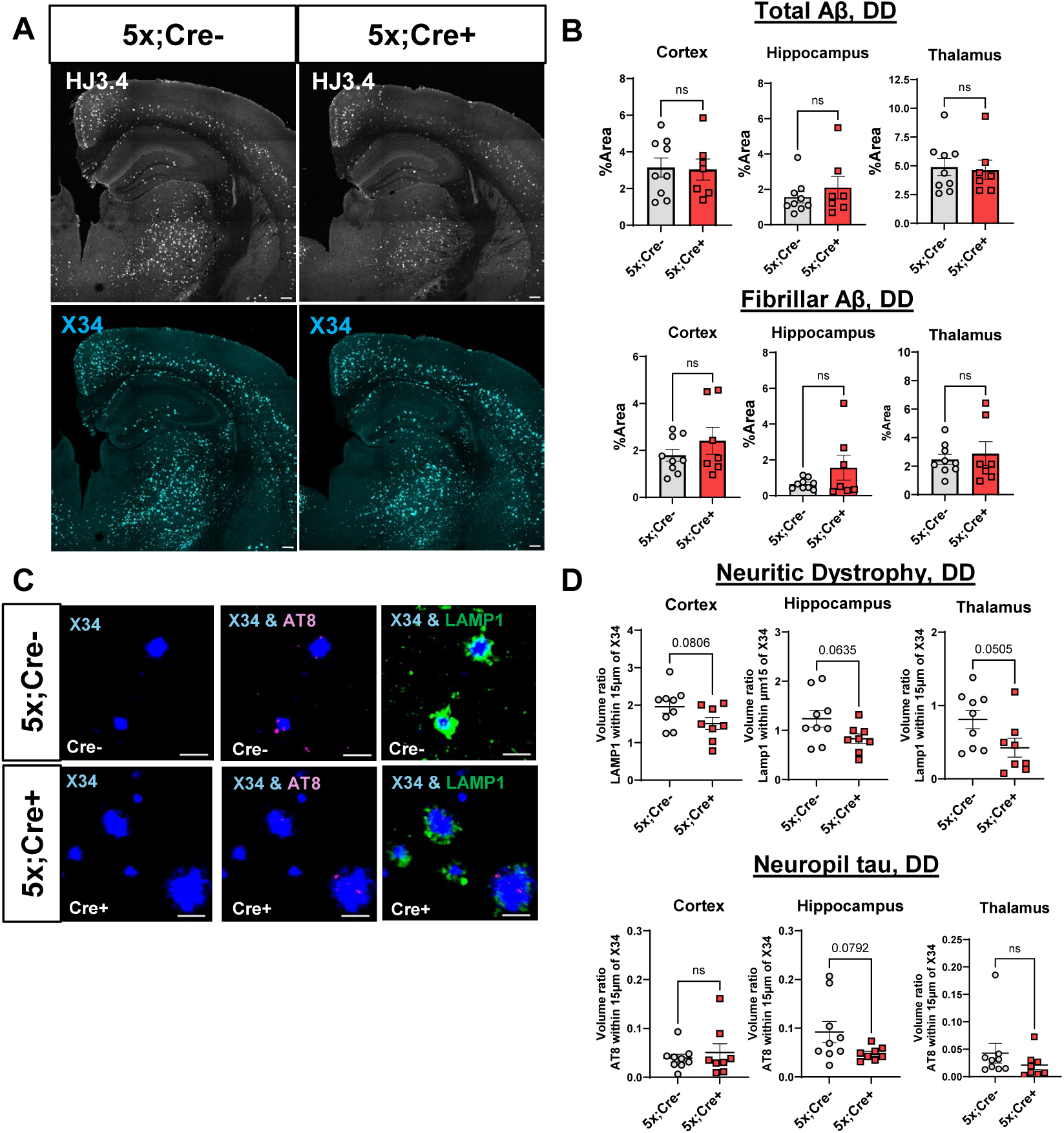
GABAergic *Bmal1* KO has minimal impact on amyloid plaque deposition and peri-plaque pathology under constant darkness conditions. **A.** Representative whole brain images from 5mo old 5x;Cre+ mice and 5x;Cre- control mice aged in constant darkness (DD), depicting staining by HJ3.4 (total plaques, white) and X34 (fibrillar plaques, cyan). (Scale bar = 200 µm) **B.** Quantification of total (top) and fibrillar (bottom) plaque aggregation from labelled regions as depicted in A. (n=7-9 mice per group as indicated. ns = not statistically significant) **C.** Representative confocal images depicting fibrillar plaque (blue, X34), phospho-tau (magenta, AT8), and neuritic dystrophy (green, Lamp1) (Scale bar, 20 µm) **D.** Quantification of AT8 and LAMP1 as depicted in (C) Top: averaged total Lamp1 volume divided by X34 volume per field. Bottom: averaged total AT8 volume divided by X34 volume. (n = 7-9 mice per group, ns= not statistically significant, p-values between 0.05 and 0.1 are displayed

Notably, we observed that 5xFAD mice kept in LD in our paradigm developed 70% more fibrillar plaques in the cortex than those kept in DD (Fig S5). This occurred in both Cre- and Cre+ mice, suggesting a Cre-independent effect of light. Similar non-significant trends were seen in the hippocampus and thalamic regions, particularly in Cre- mice.

This suggests that DD may suppress plaque formation independently of any effect of *Bmal1* deletion, and that the inhibitory effect of VGAT-BMAL1KO on plaque formation in 5xFAD mice may only be evident in the more plaque-conducive environment provided by LD.

### GABAergic *Bmal1* knockout alters APP processing in light-dark

We next sought to examine possible mechanisms mediating the effect of GABAergic *Bmal1* knockout on plaque burden, focusing specifically on LD conditions. As we did not see an effect on microglial markers, we examined amyloid precursor protein (APP) processing, as APP undergoes a series of proteolytic cleavages to be processed into Aβ. APP is first cleaved by beta secretase/BACE1 to form soluble APPβ (sAPPβ) and a β-carboxy terminal fragment (β-CTF). sAPPβ is then further cleaved by the gamma secretase complex – which includes familial AD risk gene products PSEN1 and PSEN2, to produce Aβ ^38,39^. Alternatively, APP can be processed first by alpha secretases to produce non-amyloidogenic products, including α-carboxy terminal fragment (α-CTF).

As a proxy for APP processing, we first measured the levels of mouse *Bace1* and *Psen1* transcripts in bulk hippocampus from VGAT-BMAL1KO;5xFAD mice and 5x;Cre- controls at six time-of-day timepoints. We found a statistically significant rhythm in *mPsen1* transcript in Cre- mice that was lost in 5x;Cre+ mice. The acrophase or peak of *Psen1* rhythm was at ZT20-24/0, just before the transition from the active to inactive period (Fig S6A). The overall expression of *Psen1* was also suppressed by 20% (Fig 6A). Similar examination of *Bace1* transcript revealed no rhythmic activity (Fig. S6B).

**Figure 6.**
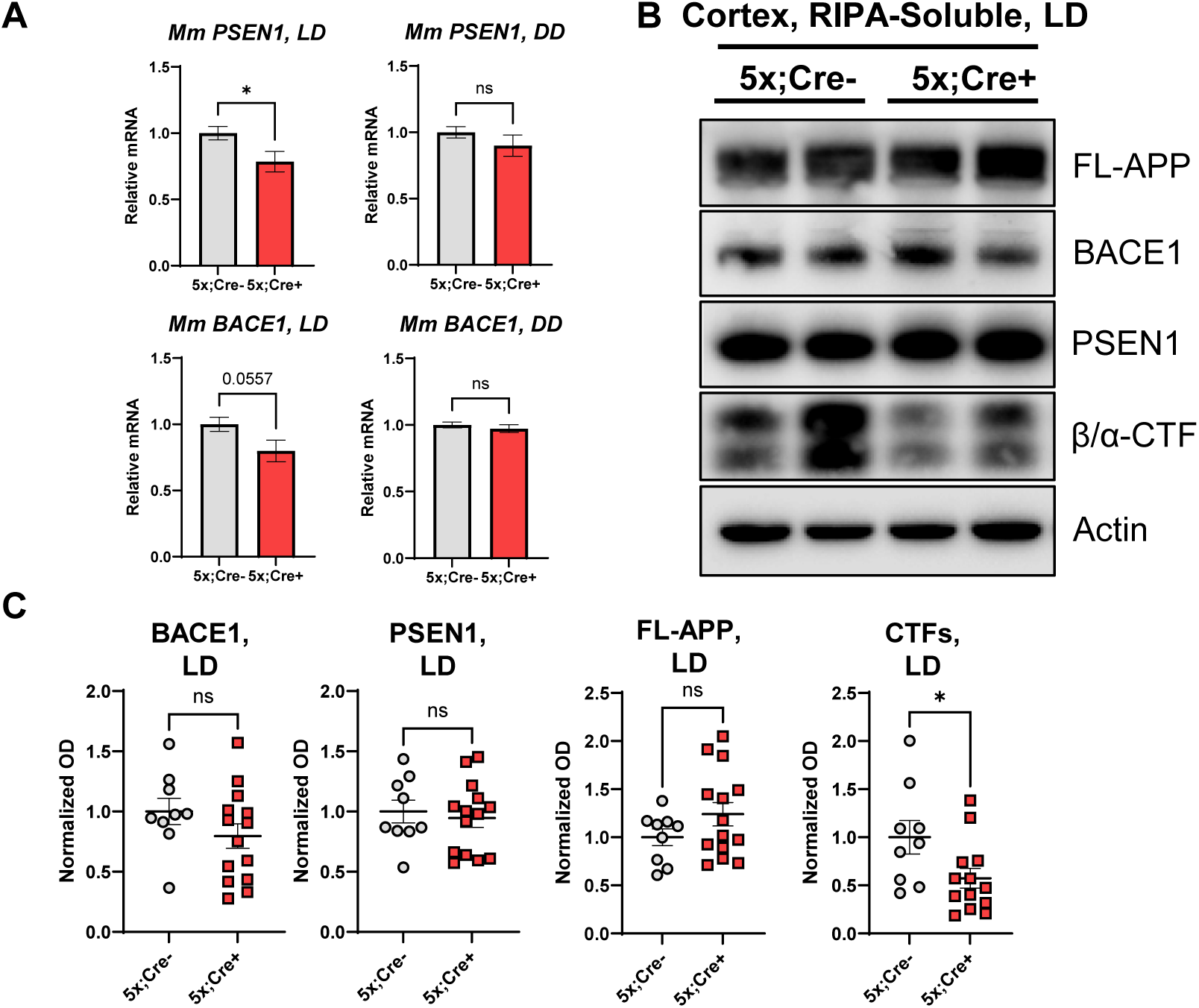
GABAergic *Bmal1* knockout alters APP processing in light-dark. **A.** Bulk expression of *Psen1* and *Bace1* transcripts from 5x;Cre+ and 5x;Cre- controls aged in LD (left) or DD (right). (n=9-15 mice per genotype per group in LD, and 6 mice per group in DD. ns=not statistically significant, * p<0.05 by Students t, p-values between 0.05 and 0.1 are displayed) **B.** Immunoblot assessing expression of APP processing enzymes (BACE1, PSEN1), full-length APP (FL-APP), and C-terminal fragments (CTF) in soluble brain lysates from cortex from 5mo 5x;Cre+ mice and 5x;Cre- control mice aged in LD. **C.** Quantification of immunoblot values as depicted in (B) (n = 9-15 mice per group as indicated, ns = not statistically significant, * p < 0.05 by Student’s t

Overall levels of *mBace1* were suppressed roughly 20%, but this difference did not quite meet statistical significance (p=0.055). Expression of both transcripts in DD mice was unchanged in Cre+ mice relative to Cre-, suggesting that suppression of mouse *Psen1* by *Bmal1*-knockout may be light-dependent (Fig 6A).

We next analyzed the levels of APP processing proteins and products via immunoblotting in cortical lysates from 5x;Cre+ mice and 5x;Cre- controls in LD. Total CTF amount (α-CTF + β-CTF) relative to full-length (FL) APP was decreased in 5x;Cre+ mice relative to Cre-; 5xFAD littermates, suggesting that total proteolytic conversion of APP via amyloidogenic and non-amyloidogenic pathways may be suppressed in VGAT- BMAL1 KO mice (Fig 6B,C). However, protein levels of BACE1 and PSEN1 protein remained similar in 5x;Cre+ mice and Cre-; 5xFAD controls (Fig. 6B,C). With regard to PSEN1, this may be partially explained by the PSEN1 antibody measuring both endogenous murine and transgenic human PSEN1 from the 5xFAD transgene, the latter of which may be unaffected GABAergic *Bmal1*-knockout. These data suggest that GABAergic *Bmal1* knockout in LD suppresses APP proteolytic processing, possibly by inhibiting expression of murine presenilin 1 transcript.

### The light-dark cycle and GABAergic *Bmal1* deletion interact to regulate the ECM transcriptome

Finally, we explored transcriptomic changes occurring in VGAT-BMAL1 KO;5xFAD mice under both LD and DD conditions. We performed bulk RNA-seq on hippocampal tissue from LD and DD-exposed 5x;Cre+ mice and 5x;Cre- controls. An initial analysis revealed 391 transcripts that were differentially regulated in Cre+ mice relative to Cre- (p < 0.005) in LD (Fig 7A). Of these, 108 were categorized as statistically significant after correction for multiple comparisons (FDR < 0.1). By contrast, under DD conditions, only 27 transcripts showed nominal differential expression between 5x;Cre- and 5x;Cre+ mice, none of which remained significant after FDR correction, underscoring the role of the light-dark cycle in gating the transcriptional response to GABAergic *Bmal1* knockout (Fig 7B). KEGG analysis of the DEG list from Cre- and Cre+;5xFAD mice in LD revealed enrichment of transcripts related to extracellular matrix (ECM) - receptor interaction (including *Col1a2, Lama5, Lamc3*) as well as lysosomal function (including *Naga, Fuca1, Glb1*) (Fig. 7C). A majority of these changes (94 out of 108) were the result of gene upregulation (Fig S7A). To identify putative upstream transcription factors that could regulate these transcripts, we performed *in silico* analysis using ChIP-X Enrichment Analysis (ChEA3), which uses a collection of libraries including ENCODE and ARCHS4 to identify transcription factor-target associations^40^. The top 10 upstream regulators by integrated mean rank were PRRX2, FOXC2, SOX18, GLIS2, FOXS1, AEBP1, ATOH8, HIC1, OSR1, and PRRX1 (Fig 7D). We then examined the rhythmicity of these factors, aside from FOXC2 and FOXS1 which were not detected, at the translatome level using our previously published mouse brain circadian gene expression atlas^41^. Of these eight factors, *Aebp1* showed rhythmicity in the largest number of conditions, including in bulk cerebral cortex tissue from both WT and APP/PS1 mice, suggesting either direct or indirect circadian regulation (Fig S7B, S7C). AEBP1, a transcritption factor involved in extracellular matrix formation, has previously been shown to be upregulated in the human AD hippocampus and associated with amyloid plaque and pre-tangle tau pathology^42,43^. We found that AEBP1 protein levels were increased by roughly 25% by western blot in cortex from VGAT-BMAL1KO;5xFAD mice (Fig 7E). Moreover, we observed that 25 known AEBP1 target genes (identified by ChEA3) were upregulated in cortex of VGAT-BMAL1KO;5xFAD mice (as compared to 5x;Cre- controls) under LD conditions, but not under DD conditions (Fig. 7F). In general, AEBP1 target gene expression was increased 40% (Fig 7F, S7D). Together, these results suggest that GABAergic *Bmal1* knockout induces extracellular matrix and lysosomal gene expression in a light-dependent manner, and implicate AEBP1 – an upstream regulator of extracellular matrix and fibrotic pathways – as a partial mediator of these transcriptomic changes^44,45^.

**Figure 7.**
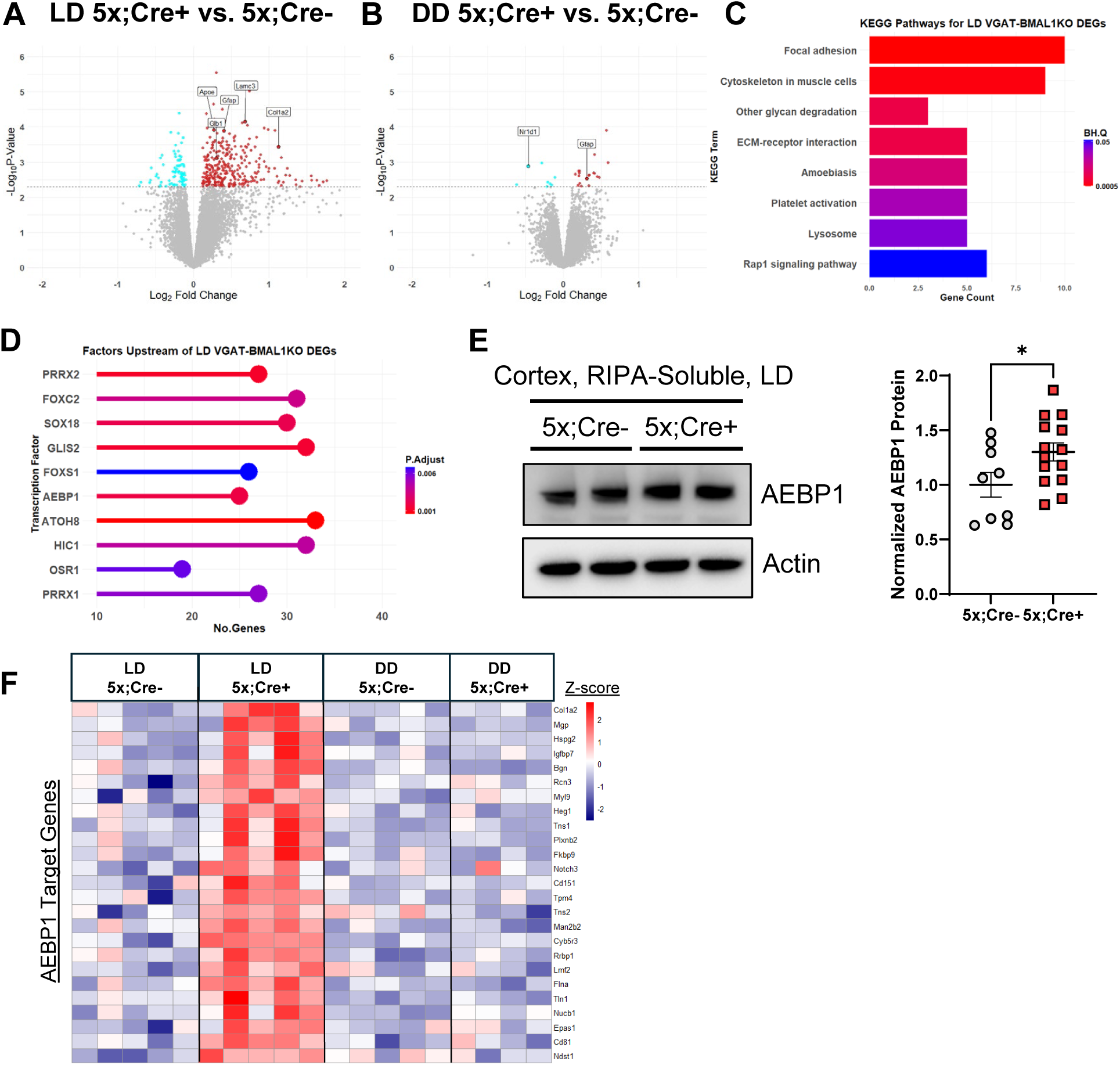
GABAergic *Bmal1* deletion promotes induction of ECM-related genes in 5xFAD mice in a light-dependent manner. **A.** Volcano plot analysis of bulk hippocampal RNAseq comparing genes from 5mo 5x;Cre+ mice and 5x;Cre- control mice aged in LD. Dashed line: p = 0.005. (n = 5 Cre-, 5 Cre+). **B.** Volcano plot analysis comparing genes from 5mo 5x;Cre+ mice and 5x;Cre- control mice aged in DD. Dashed line: p = 0.005. (n = 5 Cre-, 5 Cre+). **C.** KEGG Analysis from 5x;Cre+ vs. 5x;Cre- comparison in LD. BH.Q = Benjamin-Hochberg adjusted Q-value. **D.** Top 10 ChEA3-identified factors from 5x;Cre+ vs. 5x;Cre-, LD comparison. **E.** Quantification of AEBP1 immunoblot. (*p<0.05 by Student’s t, n = 9-15 mice per group as indicated). **F.** Heatmap of ChEA3-identified AEBP1 target genes across all conditions.

## Discussion

Disruption of circadian rhythms is highly prevalent in neurological disorders such as Alzheimer’s Disease.^46,47^ Age, the primary risk factor for developing AD, may contribute to disease progression partly through age-related circadian decline ^48^. Aβ pathology starts to develop roughly 20 years before the development of cognitive symptoms. In the vast majority of AD patients this constitutes middle age, the same epoch in which the strength of transcriptional and hormonal circadian rhythms begins to decline ^49–51^. Thus, we sought to determine whether circadian fragmentation could influence Aβ related pathology, including plaque-induced neuritic dystrophy and NP tau aggregation. Given the SCN’s principal role in regulating circadian rhythms, we employed a genetic model of SCN disruption using VGAT-iCre to drive *Bmal1* deletion within the SCN to drive. As previously reported, GABAergic *Bmal1*-knockout mice demonstrated behavioral circadian deficits in both LD and DD, as well as deconsolidated sleep and blunted clock gene rhythms within the hippocampus^17^. When crossed to 5xFAD mice, VGAT-BMAL1 KO unexpectedly mitigated amyloid plaque accumulation and plaque-associated tau phosphorylation, reduced mPSEN1 expression and amyloidogenic APP cleavage, and induced AEBP1 and ECM gene expression.

However, all of these effects only occurred when mice were kept under standard LD conditions, and not when mice were fully arrhythmic and removed from all light cues under DD conditions. Moreover, 5xFAD mice had less plaque burden in general when raised in DD rather than typical LD conditions. Our data suggest that the central circadian clock interacts with the light-dark cycle to regulate amyloid plaque accumulation and tau phosphorylation in a complex manner which was not previously appreciated.

From a general mechanistic standpoint, our experiments suggest that GABAergic *Bmal1* knockout could mitigate amyloid plaque formation by reducing Aβ production. Previous studies have shown that interstitial fluid Aβ has a strong diurnal rhythm under basal light-dark conditions in mice which is directly related to rhythms in neuronal activity^12,52,53^. However, the role of the circadian clock in regulating APP processing directly is unknown. We show that GABAergic *Bmal1* KO suppresses amyloidogenic APP cleavage to produce CTFs, suggesting reduced Aβ production. This finding is associated with loss of the normal diurnal expression rhythm of murine *PSEN1* under LD, culminating in overall decreased mPSEN1 expression. While the pool of total (human + mouse) PSEN1 protein appears to be unaffected, inhibition of the murine copy may partially explain the reduction of CTF peptides relative to full-length APP under light-dark, given that mouse PSEN1 alone is capable of cleaving human sAPPβ into plaque-generating Aβ^54^. While these data provide a partial explanation of reduced plaque load in *Bmal1* knockout mice, further studies will be necessary to examine how both light exposure and the circadian clock regulate APP metabolism at the genetic and functional levels.

Secondly, our experiments imply a previously unexplored light-dependent interaction between the GABAergic BMAL1 KO and the extracellular matrix (ECM). Structural proteins of the ECM -- including collagens and laminins -- are among the major non-Aβ components of amyloid plaques from human patients^55–57^. Biochemical experiments demonstrate that Aβ-collagen and Aβ-laminin interactions can inhibit and in some cases reverse Aβ fibril formation^58,59^. Functionally, recombinant collagen I and VI treatment can protect cultured neurons from Aβ-mediated cell death^60^. In vivo overexpression of collagen XXV in the J20 AD mouse model altered plaque aggregation in a manner similar to *Bmal1* knockout; transgenic collagen XXV overexpressing mice had reduced overall plaque load with smaller and more circular plaques^61^. While we did not manipulate the expression of these proteins directly, our results are consistent with reports that demonstrate less Aβ accumulation *in vitro* and *in vivo* after ectopic expression of ECM proteins. It is notable that one of our predicted upstream factors, AEBP1, has previously been associated with both AD pathology and connective tissue deficits in humans^42,43,45^. AEBP1 regulation of the extracellular matrix is multifaceted, as the *AEBP1* gene encodes both a transcription factor that regulates ECM gene expression and a secreted factor that activates pro-fibrotic gene expression via the Wnt/β-catenin pathway^44,62,63^. Further studies may involve directly manipulating AEBP1 expression to assess its role specifically, and that of the extracellular matrix more generally, in mediating the effect of inhibitory *Bmal1* knockout on amyloid plaque pathology.

Another finding from our study is the reduction in peri-plaque tau phosphorylation in VGAT-BMAL1 KO;5xFAD mice in LD. This was independent of the effect on overall plaque burden, as it persisted when analyzed on a per-plaque basis. This result may reflect a decrease in the toxicity of plaques in *Bmal1* knockout mice in LD and could be associated with our described changes in *Bmal1* knockout plaque morphology and/or composition. While we are unaware of any previous studies examining the effects of circadian manipulation on plaque-related tau phosphorylation, our group has previously shown that global or astrocyte-specific *Bmal1* KO can mitigate tau pathology in the P301S PS19 tauopathy model ^64^. In that work the anti-tau effect was attributed to changes in astrocyte activation and proteostatic function, while astrocytic *Bmal1* in this model is intact due to the absence of astrocytic VGAT-iCre expression. Future studies could use plaque-associated tau seeding models to examine the impact of circadian disruption on amyloid-tau interactions.

Our finding that GABAergic *Bmal1* knockout mice accumulate less amyloid and tau pathology under LD conditions is perhaps unexpected, based on previous findings from our lab, as well as the general concept that circadian disruption should be detrimental in AD^4^. We have previously shown that post-natal whole-body *Bmal1* deletion accelerated plaque accumulation in APP/PS1-21 mice, which harbor different APP and PS1 mutations than 5xFAD and tend to accumulate mostly fibrillar plaques ^65^. Thus, that study differs from this one both in terms of the amyloid model as well as the type of *Bmal1* deletion (global post-natal deletion vs. germline deletion in GABAergic neurons). BMAL1 exerts many effects on non-neuronal cell types in the brain and periphery that could impact plaque formation, while VGAT-mediated deletion should not have these effects ^66^. Future studies in which we generate mice the VGAT- BMAL1KO on the APP/PS1-21 background could help determine if these discrepancies in phenotype are due to differences between the APP/PS1-21 and 5xFAD models, which might indicate that the circadian system influences amyloid deposition differently based on the specific attributes of the model (such as the Aβ40/42 ratios, specific mutation effect, type and distribution of plaques, and rate/age of accumulation). It also remains possible that VGAT-BMAL1 KO alters neuronal inhibitory tone in the brain, leading to changes in activity-related Aβ production and aggregation (independent of effects on circadian rhythms) ^52,67,68^. However, if this were true, we would expect to see similar effects of VGAT-BMAL1 KO in both LD and DD conditions, which was not the case. Replicating our findings using a more SCN-specific Cre, or another neuron-specific Cre to delete *Bmal1* (in non-GABAergic neurons) could be an important next step.

The analysis of plaque development in both light-dark and constant darkness allowed us to evaluate the effects of GABAergic *Bmal1* knockout on Aβ pathology in both masking (LD) and non-masking (DD) environments ^69^. With respect to both circadian amplitude and period, *Bmal1* knockout mice in constant darkness exhibited a more severe circadian disruption than *Bmal1* knockout in light-dark. We expected that masking might mitigate plaque accumulation, as compared to full arrhythmicity in DD. However, Cre+ mice in LD had reduced amyloid pathology as compared to Cre- in LD, while Cre+ and Cre- mice in DD exhibited no differences in regional plaque pathology, and only non-significant reductions in neuritic dystrophy at the peri-plaque level. One possible reason for the change in plaques in light-dark but not constant darkness treated may be that *Bmal1* knockout in LD shifts circadian behavior with respect to the light cycle -- causing desynchrony between the activity rhythm and the light cycle -- as evidenced by an earlier activity onset time and *Nr1d1* gene expression (Figure 1 and Figure 2). Such a desynchrony is not possible in constant darkness, and our results suggest that this has underlying effects on Aβ pathology and related pathways that are not recapitulated by arrythmicity ^70^.

In summary, we have found that deletion of *Bmal1* specifically in VGAT-expressing GABAergic neurons, which fully disrupts central circadian clock function, reduces amyloid plaque burden and tau phosphorylation in 5xFAD, and that this effect is eliminated when mice are maintained in constant darkness. Our findings suggest complex interaction between circadian rhythms and the light-dark cycle in regulating amyloid pathology and provide new insights into how circadian function might be optimized or manipulated to prevent AD.

## Materials and Methods

### Mice

All animal procedures were approved by the Washington University IACUC and were conducted in agreement with AALAC guidelines and under supervision of the Washington University Department of Comparative Medicine. 5xFAD mice were generated from mice originally purchased through The Jackson Laboratory (MMRRC; stock no. 34848). VGAT-ires-iCre C57Bl6j (Jackson Laboratory 028862) were generated and supplied by Dr. David Weaver (Univ. of Massachusetts Chan School of Medicine, Worcester, MA, USA). VGAT-ires-iCre mice were bred at least 3 generations to generate appropriate Cre and floxed alleles, and Cre+ and Cre- littermates were compared to one another. Both male and female mice were used.

For aging 5xFAD mice in constant darkness and light-dark, mice were transferred into custom-built tents with ventilation and light manipulation. Mice were group housed, and food and water provided *ad-libitum*. We exposed mice to either a 12h:12h light/dark (LD, where lights on is defined as zeitgeber time ZT0) or 12h:12h dark/dark (DD) with temperature range 20-26 °C and 50-60% humidity. A visual inspection was performed once a week during cage changes. Unless otherwise specified, all mice were perfused and tissues harvested at 5 months of age.

### Circadian Activity Measurement

Mice were transferred into single cages in custom-built tents for no more than four weeks during recordings, and food and water provided *ad-libitum*. Mice were acclimatized for one week in LD before data acquisition. Activity was recorded via motion-tracking infrared wireless nodes (Actimetrics), one per cage. Circadian statistical measures were plotted and analyzed via Clocklab 6.1.02 and Graphpad Prism 10.4.1.

Free-running period was calculated by Χ^2^ periodogram using 8 days before the transition from LD to DD, and 8 days starting two days after the LD to DD transition. Fast Fourier Transform (FFT) was used to calculate the relative amplitude of the circadian component from 18 to 30hr. Animals with a loss of circadian rhythms (NR) were excluded from period calculations.

### Sleep Measurement

Sleep measurement was performed using the PiezoSleep mouse behavioral tracking system (Signal Solutions, LLC, Lexington, KY, USA) ^71^. This method utilized thin dielectric piezo sensors that generates a signal in response to pressure changes on its surface. Mice were individually housed in home cages with the piezo pad underneath, and were acclimated to housing conditions for one week before the start of data collection. They were given fresh bedding as well as food and water *ad libitum*. Cre- and Cre+ littermates, either 5xFAD- or 5xFAD+ were continuously monitored over the course of two weeks (7x LD, 7x DD). Data was acquired and analyzed using SleepStats software (Signal Solutions, LLC, Lexington, KY, USA).

### Immunohistochemistry

Mice were anesthetized with 150mg/kg ip pentobarbital and perfused for 3 minutes with cold Dulbecco’s modified PBS with 3% heparin. One hemisphere was dissected into cortical and hippocampal regions, flash frozen, and stored at -80 for RNA and biochemical analysis. The other was fixed in 4% PFA for 24 hours at 4°C. Brains were sectioned on a sliding microtome in 40-micron sections and stored in cryoprotectant (30% ethylene glycol, 15% sucrose 15% phosphate buffer in ddH2O). For fibrillar plaque staining, sections were washed in TBS, incubated in TBS+0.25% Triton X-100 for 30 min, then incubated in X34 staining buffer (10uM X34 and 20mM NaOH in 60% PBS, 40% ethanol) for 20 min. Sections were then blocked in 3% donkey serum for one hour, then incubated in TBSX containing 1% donkey serum and primary antibodies overnight at 4° C. The following primary antibodies were used: Rat anti-Lamp1 (DSHB, 1D4B), biotinylated AT8 (Thermo, MN1020B), goat anti-Iba1 (Abcam, ab5076), rat anti- CD68 (BioRad, MCA1957). Biotinylated HJ3.4 was generously provided by David M. Holtzman. On day 2, sections were washed and incubated in 1:1000 donkey fluorescent secondary antibodies before being mounted on slides using Prolong Gold.

### Image Acquisition and analysis

Images were obtained from 2 sections per mouse, 200 microns apart. Slides were scanned on a W1 CSU SoRa Spinning Disk Confocal (Nikon) and processed using ImageJ version 1.54 (National Institutes of Health). For whole region analysis, a stitched image was generated from a 3x by 3x tiling of 9x z-stacks (40 micron) of each section using the 10x objective generating a final image at 6682 by 6682 pixel resolution, then processed via a maximum intensity projection. Laser and detector settings were set for the acquisition of each image consistently. Peri-plaque images were acquired with a 20x objective, 2x zoom at 1,024 x 1,024 resolution.

Peri-plaque images were analyzed on a semi-automated platorm using Imaris 9.5 (Bitplane) and MATLAB. Surfaces were created using X34, dilated by 15 microns, and colocalized with various immunostained surfaces. For volume ratio calculations, the combined volume of each given surface in an image was divided by the combined X-34. All staining experiments were imaged and quantified by a blinded investigator.

### RNA Quantification and Analysis

Tissue was homogenized in 500uL of Trizol with RNAse free beads in a bullet blender for 4 minutes. Trizol samples were then extracted with 1:6 chloroform and centrifugation at 12,500g for 15 minutes. RNA was extracted from the aqueous layer using PureLink RNA Mini Kit according to the manufacturer’s protocol. Nanodrop spectrophotometry was used to determine the RNA concentration, after which the samples were converted into cDNA using a high capacity RNA-cDNA reverse transcriptase kit (Applied Biosystems) with 400-800 g RNA per 20 uL reaction. We performed qPCR using ABI Taqman Primers and ABI PCR Master Mix Buffer on ABI StepOnePLus or QuantStudio 12K thermocyclers. Taqman primers (Invitrogen) were used, and mRNA values were normalized to β-Actin (Actb) or Gapdh. For microfluidic experiments, qPCR array measurements were performed by the Washington University Genome Technology Access Center (GTAC) using a Fluidigm Biomark HD system, again using Taqman primers and normalized to GAPDH. RNA-Seq analysis was performed as previously described^72^. All gene counts were imported into the R/Bioconductor package EdgeR and TMM normalization size factors were calculated to adjust the samples for differences in library size. Genes with an average expression of more than 2 counts per million across all samples were considered as expressed. Pathway enrichment analysis was performed through clusterProfiler^73^.ChIP X-Enrichment analysis was performed using the online webtool.

### Protein Extraction

Flash frozen hippocampal brain tissue was immersed in RIPA buffer pH 8.0: 150mM NaCl, 50mM Tris, 0.5% deoxycholic acid, 1% Triton X-100, 0.1% SDS, 5mM EDTA, 20mM NaF, and 1mM sodium orthovanadate supplemented with 1% protease and phosphatase inhibitor. Tissue was ground on ice for 15 seconds using a tissue homogenizer and centrifuged at 4°C at 10,000g for 5 minutes. The resulting supernatant was taken as the soluble fraction. The pellet was incubated in 1:6 5M guanidine hydrochloride, subjected to ultrasonication for 30 seconds, and incubated at room temperature for 2 hours with shaking. The resulting homogenate was then centrifuged for at 4°C at 20,000g for 20 minutes, and the supernatant collected as the insoluble fraction for long-term storage at -80. Protein concentration was analyzed via Pierce BCA Protein Assay (Thermo).

### Western Blot

Hippocampal and cortical samples were weighed and lysed according to the protein extraction protocol specified above. Equal amounts of protein samples (10-20ug) were dissolved in sample buffer, heat treated at 95° C for 5 minutes, and treated to sodium dodecyl sulfate-polyacrylamide get electrophoresis before electrophoretic transfer to immunoblotting 0.2um PVDF membranes. The membranes were then pretreated with a blocking solution (8% dry skim milk in TBS, 0.1% Tween 20) for 1h at room temperature, then treated with primary antibodies overnight at 4° C. The following primary antibodies were used: Rabbit anti-β amyloid (Thermo, #51-2700, 1:1000), Rabbit anti-PSEN1 (CST #5643, 1:1000), Rabbit anti-BACE1 (Abcam, EPR3956, 1:1000), Rabbit anti-β actin (CST #4970), Mouse anti-AEBP/ACLP (SC Biotechnology, sc-271374). The next day, after three washes with TBST, membranes were incubated in horseradish peroxidase (HRP)-conjugated secondary antibodies against rabbit IgG in 1% BSA for 1 hour at room temperature. Proteins were detected by chemiluminescence using Super Signal Wes Pico PLUS Chemiluminescent Substrate (Thermo, #34577). Western blots were visualized using a ChemiDoc Imaging System (BioRad) and analyzed with ImageJ. Unless otherwise specified, blots were normalized to Actin.

### Statistics

Graphs display the mean+/- SEM, and n indicates the number of animals, unless otherwise indicated in the figure legend. To examine if variances were different, an F test was first performed for datasets with a single dependent variable and two groups. If not, a two-tailed unpaired t-test was performed. If variances were significantly different, a non-parametric Mann-Whitney U test was performed. For datasets with two dependent variables, 2-way ANOVA was performed. And if the main effect was significant, Sidak multiple comparisons test was applied for the variables. Outliers were identified using ROUT (Q = 1%) and excluded. Statistical tests were performed with GraphPad Prizm software version 10.4.1. P values greater than 0.1 were not as not significant (NS), while P values from 0.1-005 were specifically listed in the figure. P < 0.05 was considered significant and notes with asterisks indicating the p-value: *P < 0.05, ** < 0.005, *** < 0.0005

## Supporting information

Full Western Blot

## Declarations Availability of data

The datasets used and/or analysed during the current study are available from the corresponding author on reasonable request. RNAseq data will be available on the NIH GEO website at the time of publication.

## Competing Interests

The authors report no competing interests related to this publication.

## Funding

This work was supported by National Institute on Aging grant 5R01AG054517 (ESM), as well as the WashU Personalized Medicine Initiative (PI: Dr. David Perlmutter). MWK was supported by National Institutes of General Medical Science grant 5T32GM008151. RNA sequencing was performed by GTAC at the McDonnell Genome Institute at Washington University. AS was funded in part by the BrightFocus Foundation grant number A2024031F. The GTAC is partially supported by NCI Cancer Center Support grant P30 CA91842 to Siteman Cancer Center, ICTS.CTSA grant UL1TR002345 from the National Center for Research Resources (NCRR), a component of the NIH, and NIH Roadmap for Medical Research. Microscopy was performed at the Washington University Center for Cellular Imaging (WUCCI), which is supported by Washington University School of Medicine, The Children’s Discovery Institute of Washington University and St. Louis Children’s Hospital (CDI-CORE-2015-505 and CDI-CORE-2019-813), and the Foundation for Barnes-Jewish Hospital (nos. 3770 and 4642).

## Acknowledgements

We thank Dr. David Weaver for editorial comments on the manuscript.

## Notes

https://musieklab.shinyapps.io/Glial_Circadian_Translatome/

### Competing Interest Statement

The authors have declared no competing interest.

### Summary of Updates

RNA-Seq on the four principal mouse groups (Cre-LD, Cre+LD, Cre-DD, Cre+DD) and subsequent downstream analysis was included as figure 7; Figures 1-6 updated; Methods revised; References revised; abstract updated to include new results

