## Supplemental Fig. S1-7 for "Circadian rhythms and the light-dark cycle interact to regulate amyloid plaque accumulation and tau phosphorylation in 5xFAD mice"

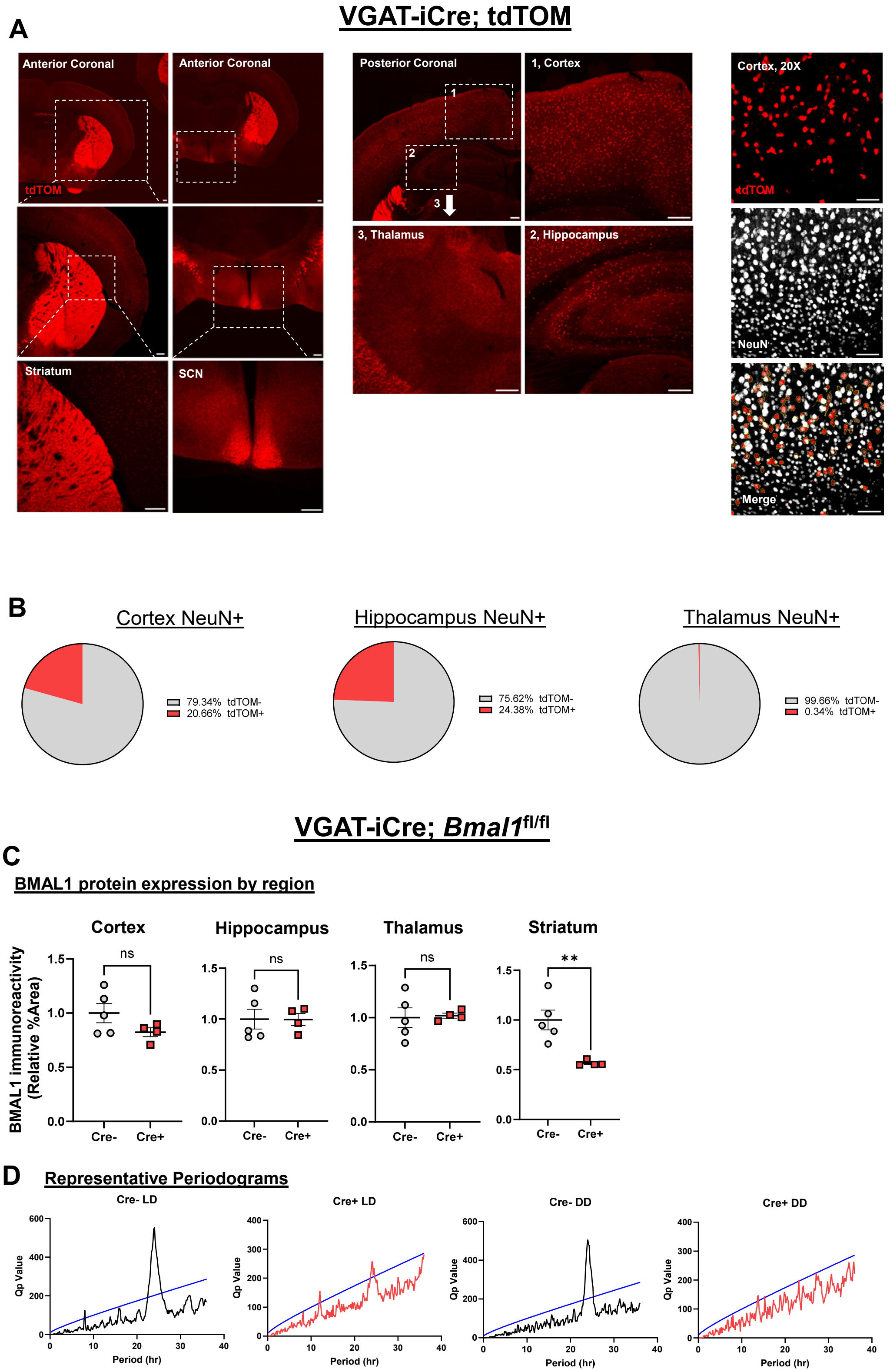


**Figure S1**

**VGAT-Cre expression and BMAL1 expression in various brain regions.**

**A.** (Top) Representative tdTOM fluorescent sections from the anterior whole brain (left), neocortex (middle) and hippocampus (left) of *VGAT-ires-iCre;tdTOM lox-stop-lox* mice. (Scale Bar = 200 μm). (Bottom) Representative 20x NeuN (white, left), tdTOM (red, middle) and merged images (right) from the neocortex. tdTOM+ cell bodies are highlighted yellow. (Scale bar = 50 μm) **B.** Pie charts of tdTOM+ nuclei among NeuN+ immunostained nuclei from *VGAT-ires-iCre;tdTOM lox-stop-lox* mice **C.** Quantification of BMAL1 immunostaining in selected regions from *VGAT-ires-iCre+;Bmal$^{fl/fl}$* mice. (n=4-5 mice per group as indicated, ns: not significant, ** Student's t, $p < 0.005$) **D.** Representative periodograms used to plot the amplitude and period data shown in figures 1E and 1F. Blue line depicts significance ($p = 0.01$), Black (Cre-) and Red (Cre+) traces depict amplitude.

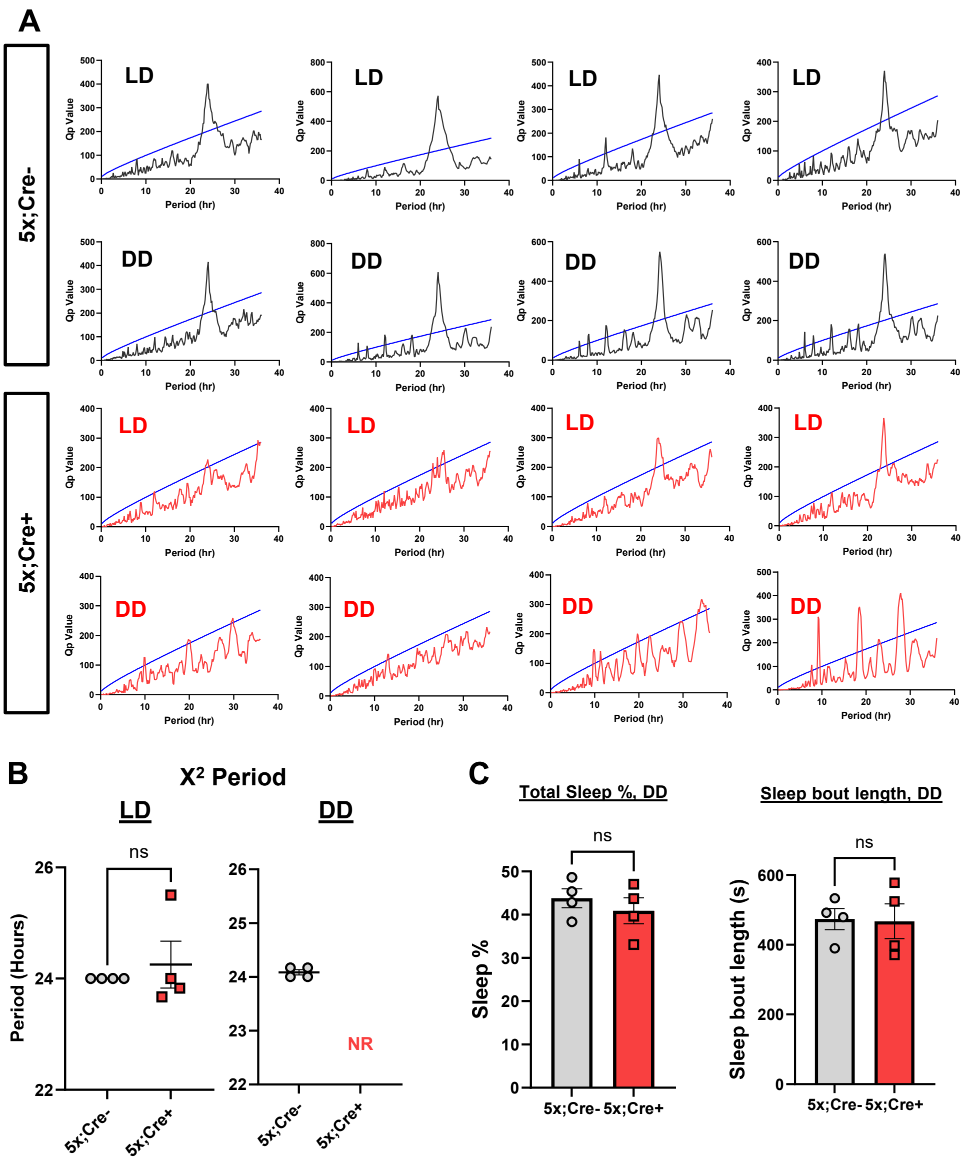


**D**

| Probe | Adj. pval (5x;Cre-) | Adj. pval (5x;Cre+) |
|---|---|---|
| *Arntl (Bmal1)* | 4.7e-6 | 0.15 |
| *Nr1d1* | 4.7e-6 | 0.002 |
| *Dbp* | 2e-6 | 0.34 |
| *Per2* | 2e-5 | 0.2 |
| *Fabp7* | 4e-4 | 1 |
| *Nr1d2* | 0.001 | 0.26 |
| *Cry1* | 0.002 | 1 |
| *Npas2* | 0.04 | 0.06 |
| *Cdkn1a* | 0.63 | 1 |

**Figure S2**

**Periodogram, transcriptional, and sleep metrics from VGAT-BMAL1KO;5xFAD mice**

**A.** LD and DD periodograms from *VGAT-ires-Cre;Bmal$^{fl/fl}$;5xFAD* (5x;Cre+) mice and 5x;Cre- controls. Blue line depicts significance (p = 0.01), Black (5x;Cre-) and Red (5x;Cre+) traces depict amplitude. **B.** $X^2$ period analysis quantification from peridograms represented in (A). No coherent rhythms (NR) were seen in Cre+ DD mice. (ns= not statistically significant) **C.** Total Sleep% and Sleep bout length in *VGAT-ires-Cre;Bmal$^{fl/fl}$;5xFAD* in DD. (ns= not statistically significant) **D.** Table of clock genes and adjusted p values as calculated by JTK_CYCLE, from Fig. 2G.

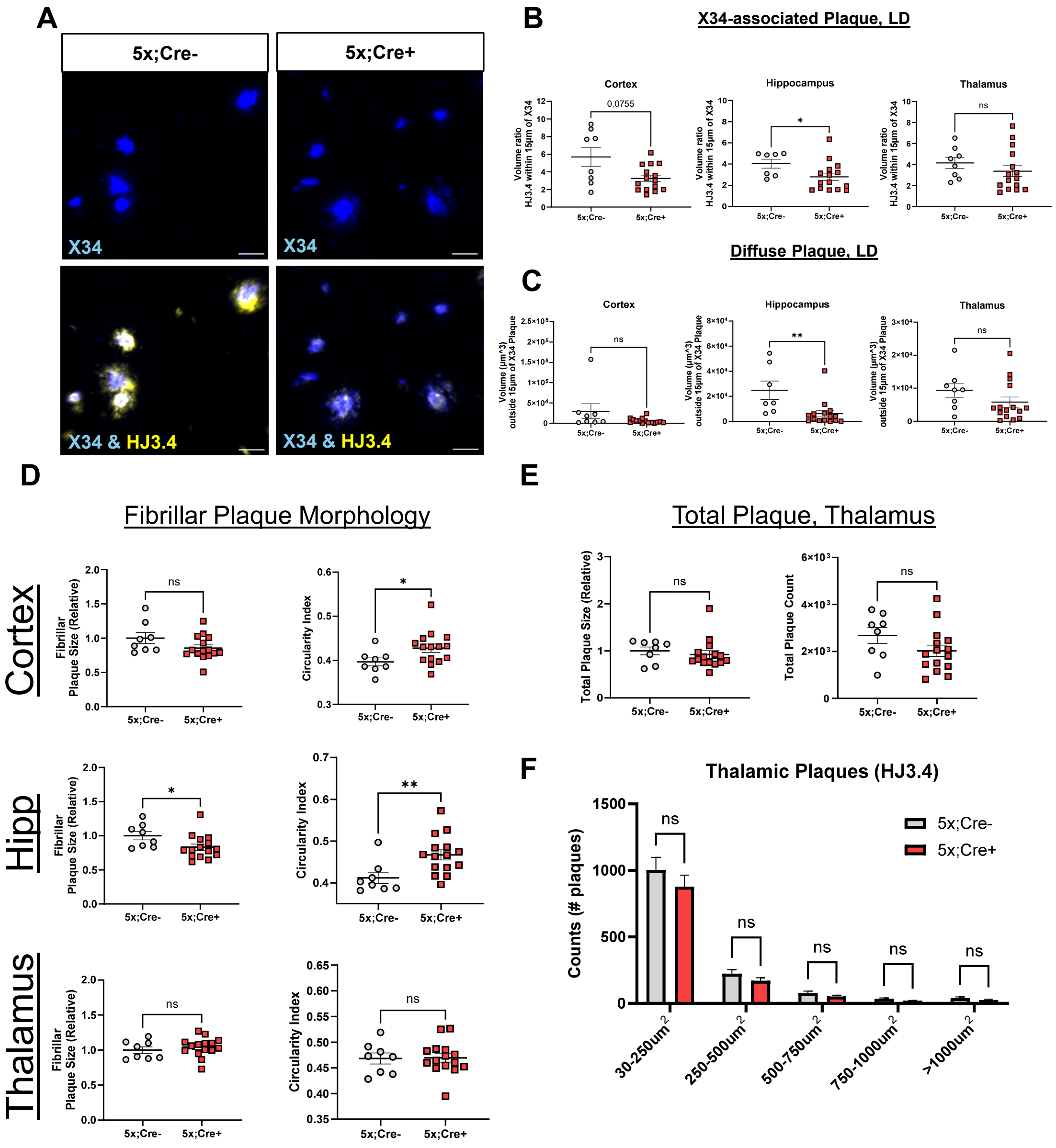


**Figure S3**

**GABAergic *Bmal1* knockout alters fibrillar plaque morphology and non-fibrillar plaque aggregation**

**A.** Representative confocal images depicting fibrillar (blue, X34) and non-fibrillar (yellow, HJ3.4) Aβ (Scale bar, 20 µm) from 5mo 5x;Cre+ mice and 5x;Cre-control mice aged in LD. **B-C** Quantification of peri-plaque HJ3.4 volume generated from 3D surfaces of MIPs depicted in A. All data depicts mean +/- SEM unless otherwise indicated. “X-34 Associated Plaque” denotes HJ3.4 volume within 15 microns of X34 volume. “Diffuse Plaque” denotes HJ3.4 volume outside 15 microns of X34 volume (n=9-15 mice per group as indicated, ns = not statistically significant, p-values between 0.05 and 0.1 are shown, *p < 0.05 by Students t). **D.** Size(first panel) and Circularity Index (second panel) of X34 plaques in selected regions (n=9-15 mice per group as indicated, ns = not statistically significant, *p <0.05 or ** p < 0.005 by Students t). **E.** HJ3.4 Plaque Size, count, and histogram distribution in for the thalamus (n=9-15 mice per group as indicated, ns = not statistically significant) F. Histogram of HJ3.4 plaque counts in thalamus. (n = 9-15 mice per group, ns = not statistically significant by Two-Way ANOVA with Šídák's multiple comparisons test)

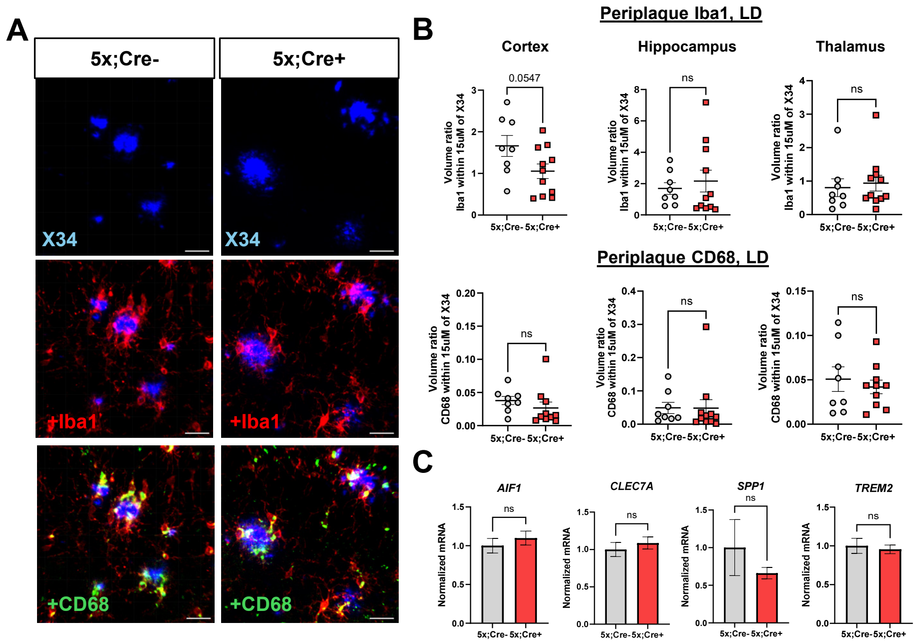


**Figure S4**

**Microglia reactivity and gene expression are unchanged in VGAT-BMAL1KO;5XFAD mice.**

**A.** Representative confocal images depicting fibrillar plaque (blue, X34), Iba1 (red) and CD68 (green) in 5mo 5x;Cre+ mice and 5x;Cre- control mice aged in LD (Scale bar, 20 µm). **B.** Quantification of staining depicted in (A) Averaged total Iba1 (top) or CD68 (bottom) volume per field divided by X34 volume. (n = 9-15 mice per group, ns=not statistically significant, p-values between 0.05 and 0.1 are displayed) **C.** Bulk hippocampal gene expression using probes against DAM-associated genes. (n=9-15 per group as indicated, ns=not statistically significant)

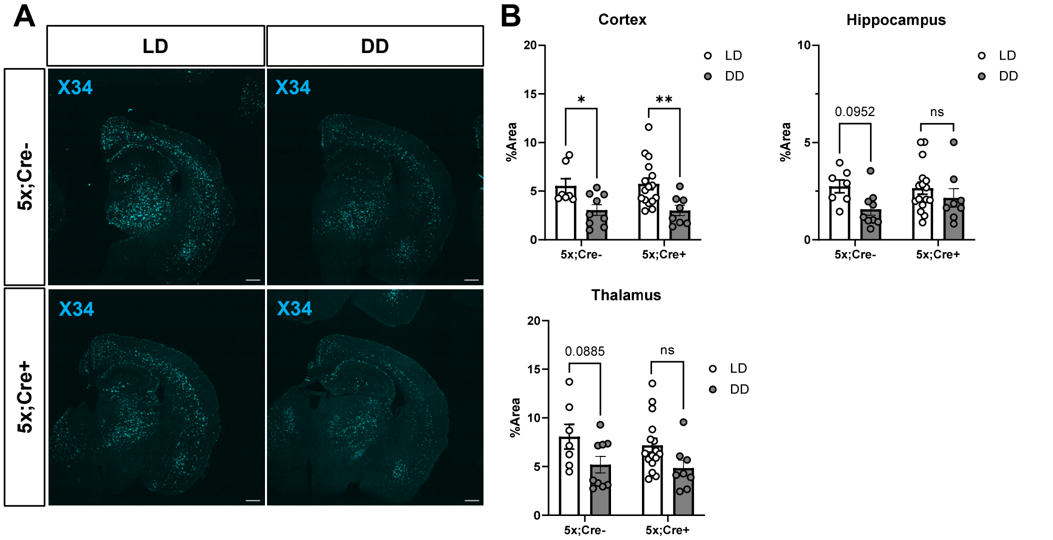


**Figure S5**

**Increased fibrillar plaque aggregation in the cortex of 5xFAD mice raised in LD as compared to DD, regardless of *Cre* genotype.**

Representative coronal brain images from 5mo 5x;Cre+ mice and Cre-; 5xFAD controls aged in LD or DD, depicting staining by X34 (fibrillar plaques, cyan). Scale bar = 500 µm **B.** Quantification of X34+ plaques in 5x;Cre+ and 5x;Cre- mice aged in LD vs DD. (n = 6-15 mice per group as indicated, *p < 0.05 or **p < 0.005 by Two Way ANOVA with Šídák's multiple comparisons test, ns = not statistically significant, p-values between 0.05 and 0.1 are displayed.)

A

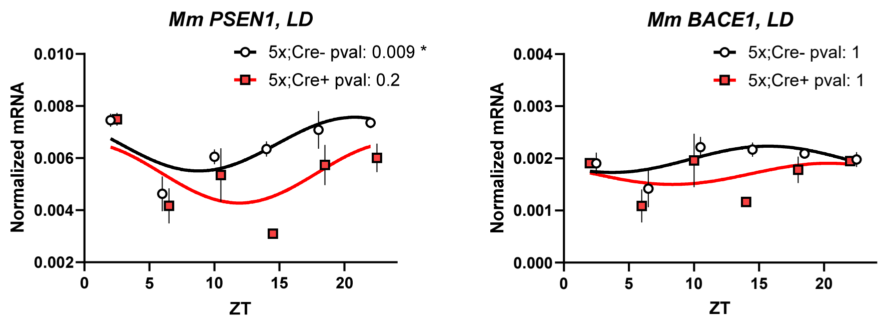


**Figure S6**

**GABAergic *Bmal1* knockout alters diurnal expression of presenilin 1**

**A. D**ata points and cosinor fit of murine *Psen1* (left) and *Bace1* (right) transcripts from selected time points in hippocampal lysates from 5mo old 5x;Cre+ and 5x;Cre- control mice aged in LD. Black circles are Cre-, Red squared are Cre+ (n=2-5 mice per genotype per time point. Benjamin-Hochberg adjusted p-values are displayed above each graph

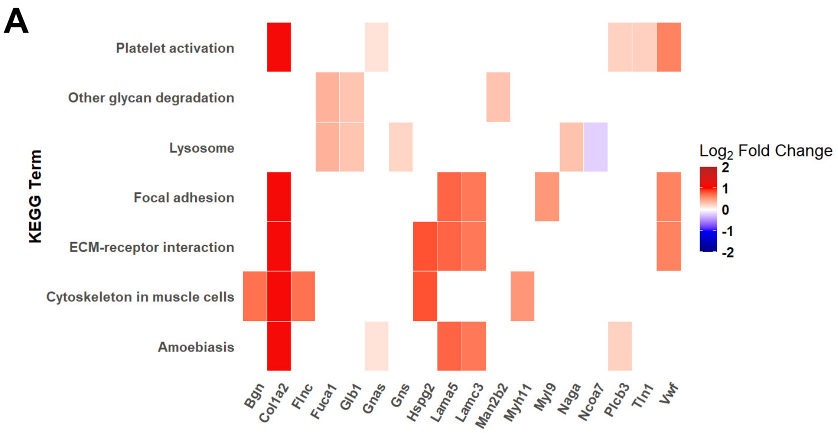


**B**

| Transcription factor | WT | APP | WT Microglia | APP Microglia | WT Astrocytes | APP Astrocytes |
|---|---|---|---|---|---|---|
| PRRX1 | | | X | | X | X |
| ATOH8 | | | | | X | |
| AEBP1 | X | X | X | | | X |
| PRRX2 | | | | | | |
| GLIS2 | | | | | | |
| SOX18 | X | X | | | | |
| HIC1 | | | X | | | |
| OSR1 | X | | | | | X |

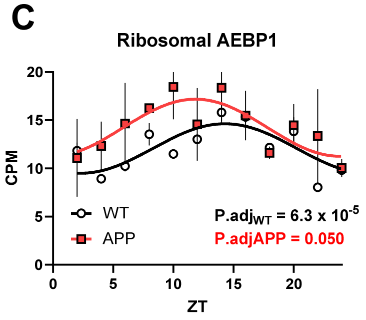


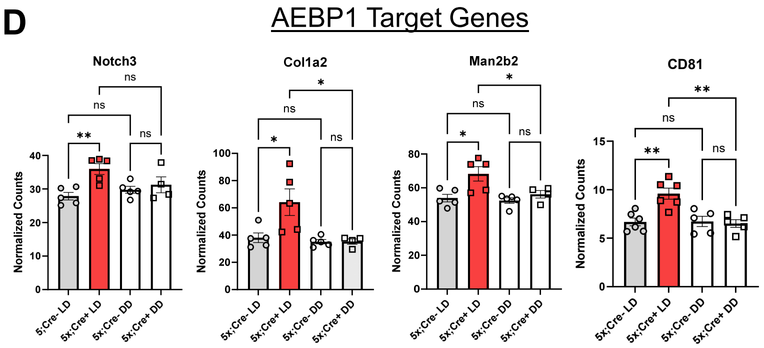


**Figure S7**

**Additional characterization of gene expression changes in 5x;Cre+ and 5x;Cre- mice**

**A.** Heatplot of KEGG terms and associated core enrichment genes from 5x;Cre+ vs. 5x;Cre- mice in LD, limited to the 5 most upregulated or downregulated genes in each term **B.** Translatome rhythmicity of ChEA3-identified transcription factors from Fig. 7D, based on data from Sheehan et al[41]. **C.** Circadian *Aebp1* transcript expression in bulk cortex from 5mo wild-type and APP/PS1-21 AD mice, based on data from Sheehan et al.[41] **D.** Selected AEBP1 target gene expression from 5x;Cre- and 5x;Cre+ mice in LD and DD (ns=not significant, *p<0.05, **p<0.005 by Two Way ANOVA with Šídák's multiple comparisons test, n=4-5 mice per group as indicated).
