## Supplementary figures and images for "Circadian rhythms and the light-dark cycle interact to regulate amyloid plaque accumulation and tau phosphorylation in 5xFAD mice"

### Full Western Blot

#### Blot 1

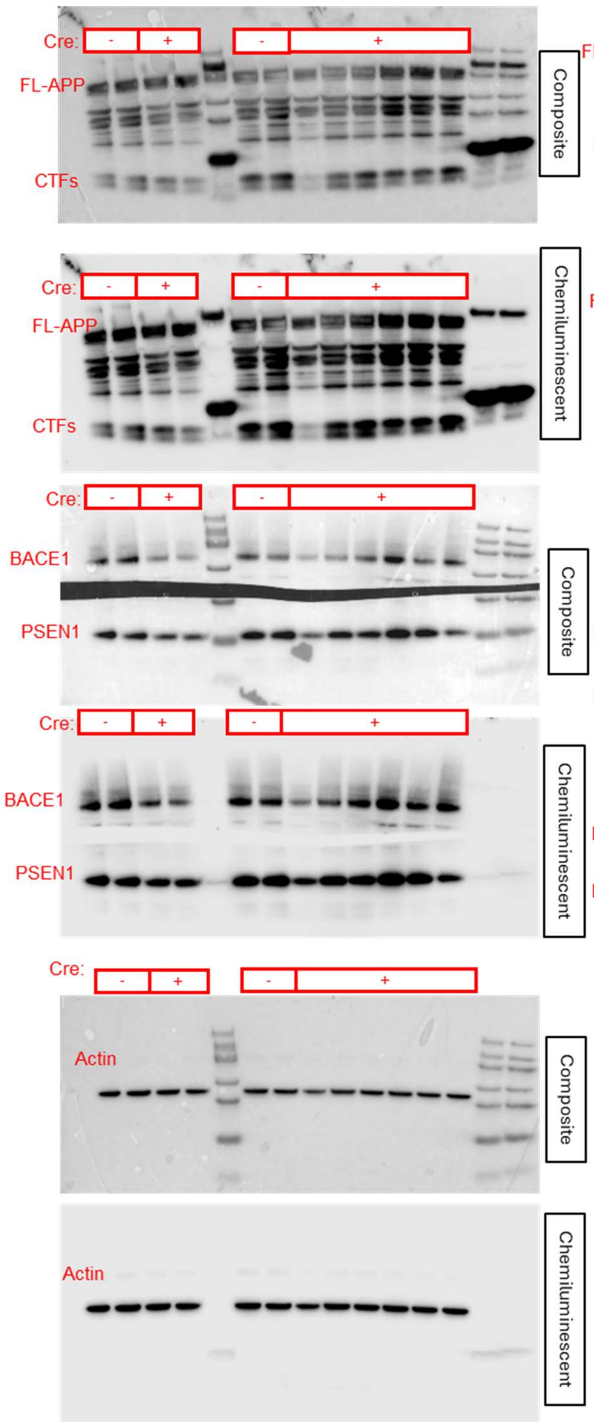


#### Blot 2

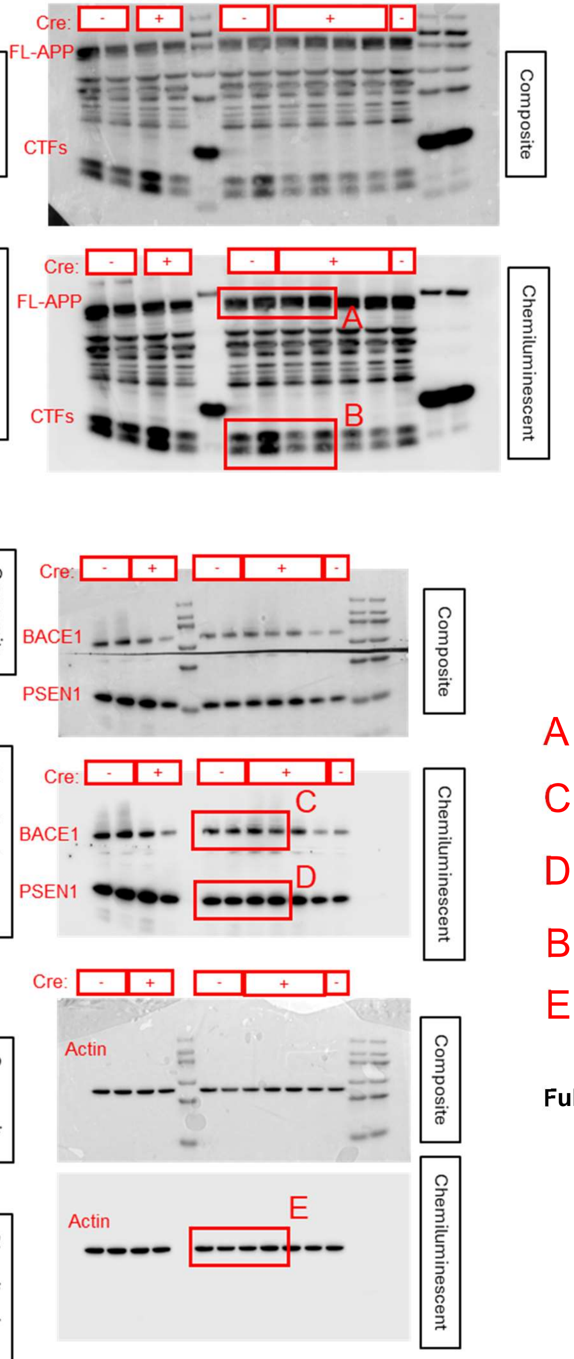


#### Figure 6B

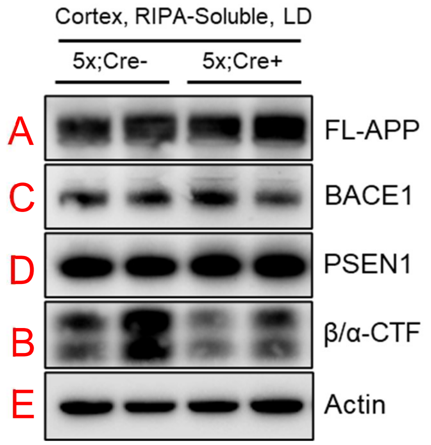


**Full immunoblot depicted in Figure 6**

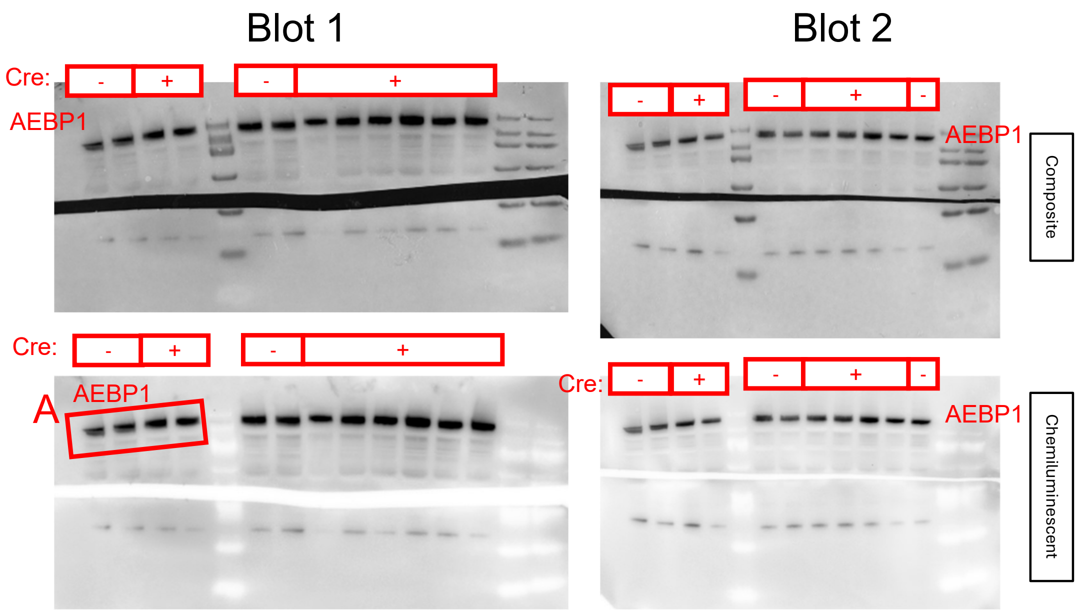


Actin reproduced from 6S

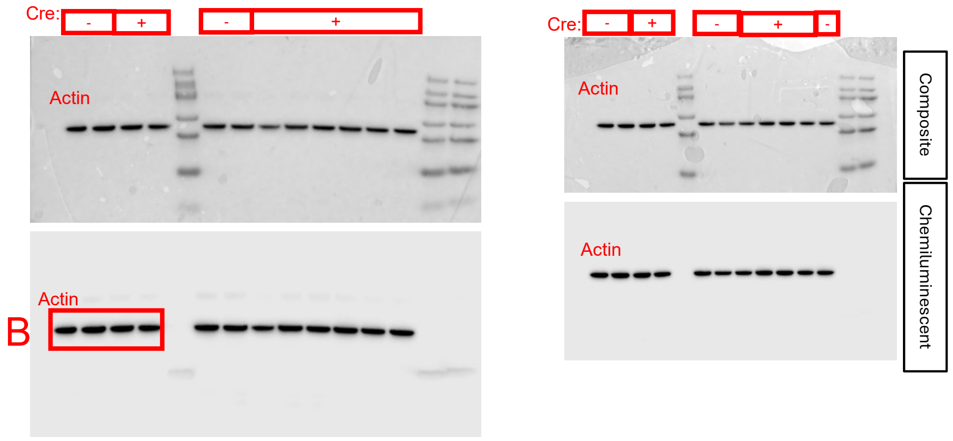


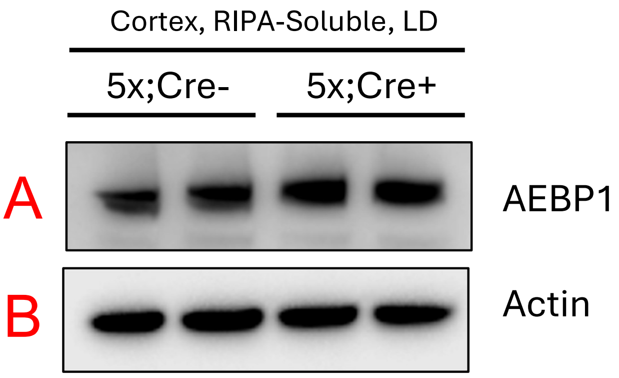
